# p38α-MAPK-deficient myeloid cells ameliorate symptoms and pathology of APP-transgenic AD mice

**DOI:** 10.1101/2021.10.18.464870

**Authors:** Qinghua Luo, Laura Schnöder, Wenlin Hao, Kathrin Litzenburger, Yann Decker, Inge Tomic, Michael D. Menger, Klaus Fassbender, Yang Liu

## Abstract

Microglial activation is a hall marker of Alzheimer’s disease (AD); its pathogenic role and regulating mechanisms are unclear. p38α-MAPK, a stress-responding kinase, is activated in AD brain in early disease stages. In APP-transgenic mice, we deleted p38α-MAPK in whole myeloid cells from birth or specifically in microglia from 9 months, and analysed AD pathology at the age of 4, 9 and 12 months. In both experimental settings, p38α-MAPK deficiency decreased cerebral Aβ and improved cognitive function of AD mice; however, p38α-MAPK-deficient myeloid cells were more effective than p38α-MAPK-deficient microglia in preventing AD pathogenesis. Deficiency of p38α-MAPK in myeloid cells inhibited the inflammatory activation of individual microglia by 4 months, but enhanced it by 9 months. Inflammatory activation was essential for p38α-MAPK deficiency to promote microglial internalization of Aβ. Interestingly, p38α-MAPK deficiency in peripheral myeloid cells reduced *il-17a* transcription in CD4-positive spleen cells. By cross-breeding APP-transgenic mice and IL-17a knockout mice, we further observed that IL-17a deficiency activated microglia and decreased Aβ deposits in AD mouse brain. Thus, p38α-MAPK deficiency in myeloid cells prevents AD pathogenesis, perhaps through reducing IL-17a-expressing T lymphocytes, and promoting Aβ clearance in the brain. Our study supports p38α-MAPK as a novel target for AD therapy.

## Introduction

Alzheimer’s disease (AD) is pathologically characterized by three components: (i) extracellular deposits of amyloid β peptide (Aβ), (ii) intracellular neurofibrillary tangles (NFT) primarily composed of hyper-phosphorylated tau protein (p-tau), and (iii) microglia-dominated inflammatory activation in the brain parenchyma (Querfurth and LaFerla, 2010). Interactions between Aβ, p-tau and inflammatory activation are primarily responsible for the progressive neurodegeneration in AD. Aβ aggregates injure synaptic junctions and neurons not only directly (Shankar et al., 2008), but also indirectly by triggering neurotoxic inflammatory activation (Wyss-Coray and Rogers, 2012). Both Aβ and inflammatory activation induce tau phosphorylation (Ghosh et al., 2013; Ryan et al., 2009), triggering the spread of NFT from the medial temporal cortex to the entire neocortex (Maphis et al., 2015; Pascoal et al., 2021; Shimada et al., 2017). Aggregation of p-tau further induces microglial inflammatory activation (Yoshiyama et al., 2007). However, activated microglia also exert a protective effect on neurons in AD by taking up Aβ peptides (Michaud et al., 2013) and promoting p-tau degradation in neurons (Qin et al., 2016).

The pathogenic role of microglial activation in AD has been extensively investigated (Heneka et al., 2015), but the microglial effects on Aβ pathology and neuronal degeneration in AD are inconclusive. For example, it is known that rare variants in the triggering receptor expressed on myeloid cells-2 (TREM2) gene increase the risk of developing AD (Carmona et al., 2018). However, one group reported that TREM2 deficiency in APP-transgenic mice increases hippocampal Aβ burden and accelerates neuron loss (Wang et al., 2015), while another group showed that TREM2 deletion reduces cerebral Aβ accumulation (Jay et al., 2015). Subsequent work suggested that the effect of TREM2 deficiency on cerebral Aβ accumulation depends on the stage of disease (Jay et al., 2017; Parhizkar et al., 2019). A recent study indicated that knockdown of TREM2 expression at a post-plaque, but not pre-plaque, or early plaque stage, reduces Aβ deposits in APP-transgenic mice (Schoch et al., 2021). Consistent with this conclusion is the observation from a longitudinal imaging study of human subjects with mild cognitive impairment (MCI) that several peaks of microglial activation appear over the disease trajectory (Fan et al., 2017). These studies underscore the effects of the changing cellular environment and reinforce the idea that the pathogenic role of microglial activation should be dynamically investigated during disease progression.

It is generally accepted that microglia play a primary role in AD pathogenesis (Reed-Geaghan et al., 2020). However, AD is a systemic disease involving peripheral immune dysregulation (Croese et al., 2021). C-C chemokine receptor type 2 (CCR2) is expressed on peripheral myeloid cells but not on microglia; however, knockout of CCR2 in a mouse model of AD resulted in fewer Aβ-associated myeloid cells and accelerated disease progression (El Khoury et al., 2007). In another study, adoptive transfer of bone marrow-derived monocytes, which express macrophage colony-stimulating factor receptor, reduced cerebral Aβ load and improved cognitive function of APP-transgenic mice (Koronyo-Hamaoui et al., 2020). Peripheral myeloid cells might also regulate T cell differentiation, which in turn affects the microglial activation. CD4-positive T cells are essential for microglial maturation (Pasciuto et al., 2020). Transient depletion of T regulatory (Treg) lymphocytes in blood significantly changes microglial responses and Aβ deposits (Baruch et al., 2015; Dansokho et al., 2016), although the two research groups had conflicting results concerning the effect of Treg lymphocyte depletion on Aβ levels.

p38α mitogen-activated protein kinase (p38α-MAPK) is a protein kinase present in a variety of cells that respond to external stress stimuli (Kumar et al., 2003). It is activated in both neurons and microglia in brains of AD patients and animal models (Hensley et al., 1999; Maphis et al., 2015; Sun et al., 2003). Our recent study indicates that p38α-MAPK deficiency in neurons reduces both Aβ and p-tau levels in the brain of AD mice (Schnöder et al., 2020; Schnöder et al., 2016; Schnöder et al., 2021). We believe that p38α-MAPK could be an effective target for AD therapy. However, the pathogenic role of microglial p38α-MAPK during the disease progress remains unclear. Systemic administration of chemical p38α-MAPK inhibitors has been shown to reduce inflammatory activation in the brain of APP- or tau-transgenic mice and improve their cognitive function (Bachstetter et al., 2012; Maphis et al., 2016). However, pharmacological treatments with p38α-MAPK inhibitors affect both microglial p38α-MAPK and neuronal p38α-MAPK, without the ability to distinguish their effects. A recent phase 2 clinical trial showed that a 24-week treatment with p38α-MAPK inhibitor decreased tau proteins in the cerebral spinal fluid of mild AD patients; however, did not improve the cognitive function (Prins et al., 2021). It was necessary to further decipher the detailed pathogenic mechanism of p38α-MAPK, which provides information to optimize AD therapy with p38α-MAPK inhibitors.

In this study, we conditionally knocked out *mapk14* gene (encoding p38α-MAPK) in the myeloid cell lineage or specifically in microglia in APP-transgenic mice and investigated the AD pathology and microglial activation in the early and late disease stages. We observed that deletion of p38α-MAPK attenuated Aβ load and neuronal deficits of AD mice; however, the pathogenic role of microglial p38α-MAPK is evolving during the disease progresses, which is potentially regulated by the peripheral interleukin (IL)-17a-expressing T lymphocytes.

## Results

### Establishment of APP-transgenic mice deficient of p38α-MAPK in myeloid cells

To investigate the pathogenic role of p38α-MAPK in microglia and peripheral myeloid cells in AD, we cross-bred APP-transgenic mice with *mapk14*-floxed and LysM-Cre^+/-^ mice, as we did in a previous study (Liu et al., 2014), to obtain APP^tg^p38^fl/fl^LysM-Cre^+/-^ (p38α deficient) and APP^tg^p38^fl/fl^LysM-Cre^-/-^ (p38α wild-type) of genotypes. By detecting *mapk14* gene transcripts in CD11b^+^ cells isolated from 9-month-old APP-transgenic mice, we estimated the rate of LysM-Cre-mediated *mapk14* gene recombination in microglia to be ∼ 45% (*mapk14*/*gapdh*: 0.146 ± 0.013 and 0.261 ± 0.046, in APP^tg^p38^fl/fl^LysM-Cre^+/-^ and APP^tg^p38^fl/fl^LysM-Cre^-/-^ mice, respectively; *t* test, *p* = 0.032), which was consistent with previous observations (∼ 40%) (Goldmann et al., 2013; Liu et al., 2014). The efficiency of LysM-Cre-mediated recombination of *mapk14* gene in circulating monocytes should be > 60% (Goldmann et al., 2013; Liu et al., 2014). To examine the recruitment of peripheral myeloid cells into the brain, APP^tg^p38^fl/fl^LysM-Cre^+/-^ mice were cross-bred with CCR2-RFP reporter mice that express red fluorescent protein (RFP) under the control of CCR2 promoter (Saederup et al., 2010). Very few RFP-positive cells were detected in close proximity to blood vessels in AD mouse brain (see supplementary Fig. 1, A). Flow cytometric analysis showed that there were 2.54% ± 0.20 % and 2.96% ± 0.22% of CD11b+ cells expressing RFP in the brains of 9-month-old p38α-MAPK-deficient and wildtype APP-transgenic mice, respectively (see supplementary Fig. 1, C and D). As LysM-Cre was reported to mediate gene recombination in neurons (Orthgiess et al., 2016), APP^tg^p38^fl/fl^LysM-Cre^+/-^ mice were further mated to ROSA^mT/mG^ Cre reporter mice, which express GFP in Cre-expressing cells (Muzumdar et al., 2007). In 6-month-old APP-transgenic mice, we detected most GFP-positive cells co-stained by antibodies against ionized calcium-binding adapter molecule (Iba)-1, a marker for microglia/macrophages, and rarely co-stained by antibodies against the neuronal marker, NeuN (see supplementary Fig. 2), which was in line with the results from our and other groups’ studies (Goulielmaki et al., 2020; Liu et al., 2019; Liu et al., 2014).

### Deletion of p38α-MAPK in myeloid cells improved the cognitive function of APP-transgenic mice

We used the Morris water maze test to examine cognitive function of mice. During the acquisition phase, 9-month-old non-APP-transgenic (APP^wt^) littermate mice with or without deletion of p38α-MAPK in myeloid cells (APP^wt^p38α^fl/fl^LysM-Cre^+/-^ and APP^wt^p38α^fl/fl^LysM-Cre^-/-^) showed no significant differences in either swimming time or swimming distance before climbing onto the escape platform (Fig. 1, A and B; two-way ANOVA, *p* > 0.05). Compared to APP^wt^p38α^fl/fl^LysM-Cre^-/-^ littermates, 9-month-old APP^tg^p38α^fl/fl^LysM-Cre^-/-^ mice with normal p38α-MAPK expression spent significantly more time (Fig. 1, A; two-way ANOVA, *p* < 0.05) and traveled longer distances (Fig. 1, B; two-way ANOVA, *p* < 0.05) to reach the escape platform. Interestingly, APP^tg^p38α^fl/fl^LysM-Cre^+/-^ mice with the deletion of myeloid p38α-MAPK performed significantly better than their APP^tg^p38α^fl/fl^LysM-Cre^-/-^ littermates in searching and finding the platform after 3 days of training (Fig. 1, A and B; two-way ANOVA, *p* < 0.05). The swimming velocity did not differ between p38α-MAPK-deficient and wild-type APP-transgenic mice or for the same mice on different training dates (Fig. 1, C; two-way ANOVA, *p* > 0.05).

**Fig. 1.**
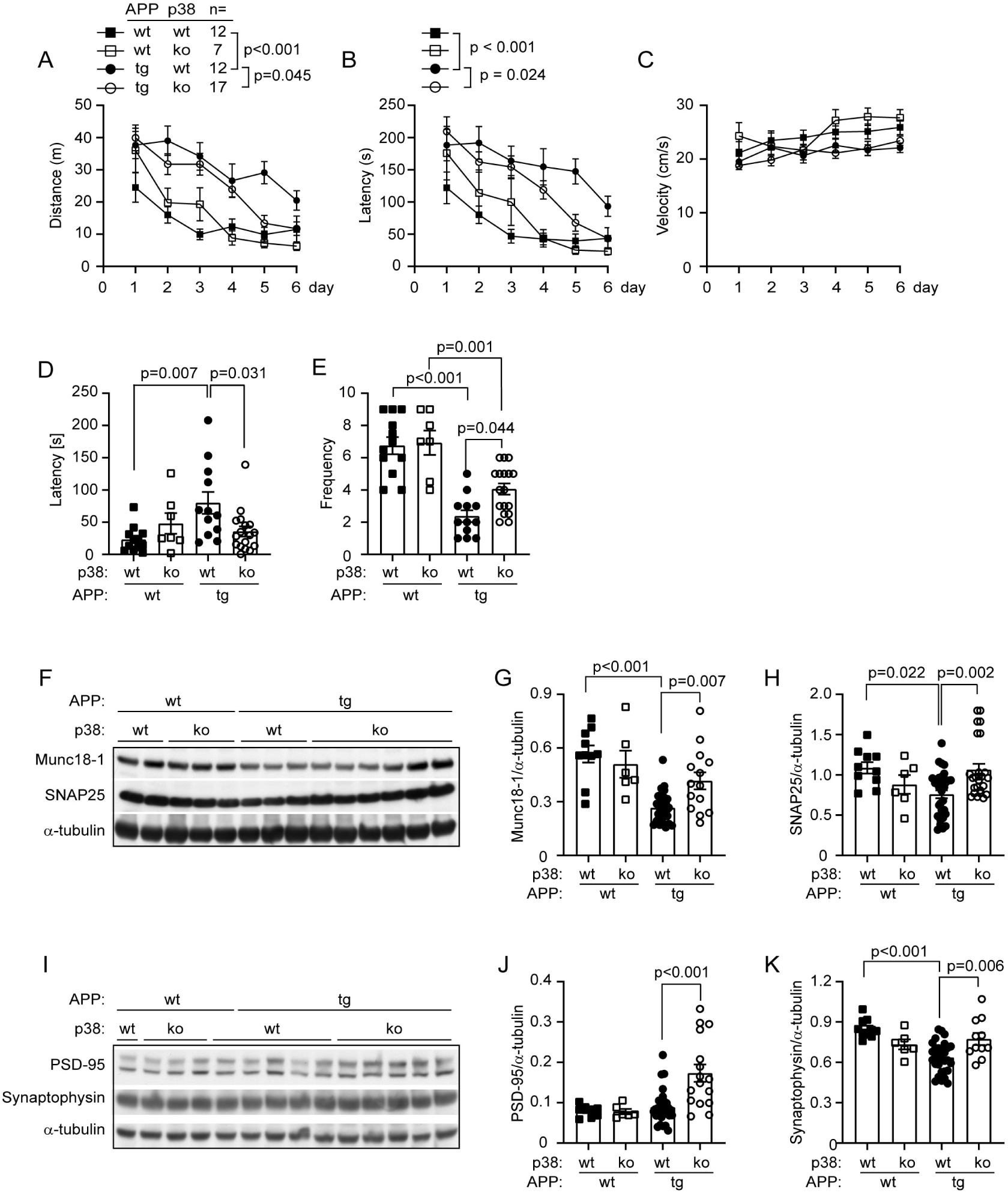
Deletion of p38α-MAPK in myeloid cells improves cognitive function and attenuates AD-associated loss of synaptic proteins in APP-transgenic mice. During the training phase of the water maze test, 9-month-old APP-transgenic mice (APPtg) spent more time and traveled longer distances to reach the escape platform than did their non-APP-transgenic littermates (APPwt). Compared to mice with normal expression of p38α-MAPK (p38α wt), deletion of p38α-MAPK in myeloid cells (p38α ko) significantly reduced the traveling time and distance of APPtg mice but not of APPwt mice after 3 days of training (A, B; two-way ANOVA from day 3 to day 6 followed by Bonferroni *post hoc* test; *n* is shown in the figure). However, deficiency of myeloid p38α-MAPK affected the swimming speed neither of APPtg and APPwt mice, nor for each mouse at different training time points (C; two-way ANOVA, *p* > 0.05). In the probe trial, APPtg mice spent significantly longer time in the first visit to the region where the platform was previously located, and crossed the platform region with significantly less frequency during the total 5-minute experiment; deletion of p38α-MAPK in myeloid cells partially recovered these APP expression-induced cognitive impairments (D, E; one-way ANOVA followed by Bonferroni *post hoc* test). Western blotting was used to detect the amount of synaptic structure proteins, Munc18-1, SNAP25, synaptophysin, and PSD-95 in the brain homogenate of 9-month-old APPtg and APPwt mice (F - K). Overexpression of APP significantly decreased proteins of Munc18-1, SNAP25, and synaptophysin in the mouse brain. Deficiency in myeloid p38α-MAPK was associated with higher levels of all these 4 tested proteins in the APPtg mouse, but not in the APPwt mouse (one-way ANOVA followed by Bonferroni *post hoc* test; *n* ≥ 11 per group for APPtg mice and 6 per group for APPwt mice).

Twenty-four hours after the end of training phase, the escape platform was removed and a 5-minute probe trial was performed to test the memory of mice. Compared to APP^wt^p38α^fl/fl^LysM-Cre^-/-^ littermates, APP^tg^p38α^fl/fl^LysM-Cre^-/-^ mice remained for a significantly longer time in their first visit to the region where the platform had been located, and crossed the original platform region with significantly less frequency during the total 5-minute probe trial (Fig. 1, D and E; one-way ANOVA, *p* < 0.01). Interestingly, when compared to APP^tg^p38α^fl/fl^LysM-Cre^-/-^ mice, APP^tg^p38α^fl/fl^LysM-Cre^+/-^ mice were able to reach the original platform region in significantly less time and crossed the region more frequently (Fig. 1, D and E; one-way ANOVA, *p* = 0.031 and 0.044, respectively). We observed differences in neither parameter analyzed in the probe trial between APP^wt^p38α^fl/fl^LysM-Cre^+/-^ and APP^wt^p38α^fl/fl^LysM-Cre^-/-^ littermate mice (Fig. 1, D and E; one-way ANOVA, *p* > 0.05).

We further used Western blot analysis to quantify the levels of four synaptic-structure proteins: Munc18-1 protein mammalian homolog (Munc18-1), synaptophysin, synaptosome-associated protein 25 (SNAP-25), and postsynaptic density protein 95 (PSD-95) in the brain homogenate of 9-month-old APP^tg^ and APP^wt^ littermate mice. As shown in Fig. 1, F-K, protein levels of Munc18-1, synaptophysin and SNAP-25 in APP^tg^p38α^fl/fl^LysM-Cre^-/-^ mice were significantly lower than levels of these proteins derived from APP^wt^p38α^fl/fl^LysM-Cre^-/-^ littermate mice (one-way ANOVA, *p* < 0.05). The reduction in Munc18-1, synaptophysin and SNAP-25 proteins due to APP-transgenic expression was rescued by the deletion of p38α-MAPK in myeloid cells (Fig. 1, G, H and K; one-way ANOVA, *p* < 0.05). PSD-95 protein levels were significantly higher in brains from APP^tg^p38α^fl/fl^LysM-Cre^+/-^ mice than that from APP^tg^p38α^fl/fl^LysM-Cre^-/-^ control mice (Fig. 1, J; one-way ANOVA, *p* < 0.001). Comparison of APP^wt^p38α^fl/fl^LysM-Cre^+/-^ and APP^wt^p38α^fl/fl^LysM-Cre^-/-^littermate mice showed no significant differences in protein levels of these four tested synaptic proteins (Fig. 1, F-K; one-way ANOVA, *p* > 0.05).

### Deletion of p38α-MAPK in myeloid cells reduces Aβ load in the brain of APP-transgenic mice

After observing that deletion of myeloid p38α-MAPK attenuates the cognitive deficits in APP^tg^ mice but not in APP^wt^ littermates, we analyzed the effects of myeloid p38α-MAPK on Aβ pathology in the APP^tg^ mice, as Aβ is the key molecule leading to neurodegeneration in AD (Mucke and Selkoe, 2012). We used standard immunohistological and stereological *Cavalieri* methods to measure Aβ volume, adjusted relative to the volume of analyzed tissues, in 9-month-old APP^tg^p38α^fl/fl^LysM-Cre^+/-^ and APP^tg^p38α^fl/fl^LysM-Cre^-/-^ mice. The volume of immunoreactive Aβ load in APP^tg^p38α^fl/fl^LysM-Cre^+/-^ mice (3.25 % ± 0.38 % in the hippocampus and 2.83 % ± 0.28 % in the cortex) was significantly lower than that in APP^tg^p38α^fl/fl^Cre^-/-^ mice (4.83 % ± 0.57 % in the hippocampus and 4.63 % ± 0.31 % in the cortex; Fig. 2, A and B; *t* test, *p* < 0.001 and = 0.039, respectively). The brain tissue was also stained with Congo red that typically binds to the β sheet structure of Aβ plaques, which showed that deficiency of p38α-MAPK decreased the cerebral level of Aβ aggregates (Fig. 2, C and D; *t* test, *p* < 0.05), corroborating the results from immunohistochemistry.

**Fig. 2.**
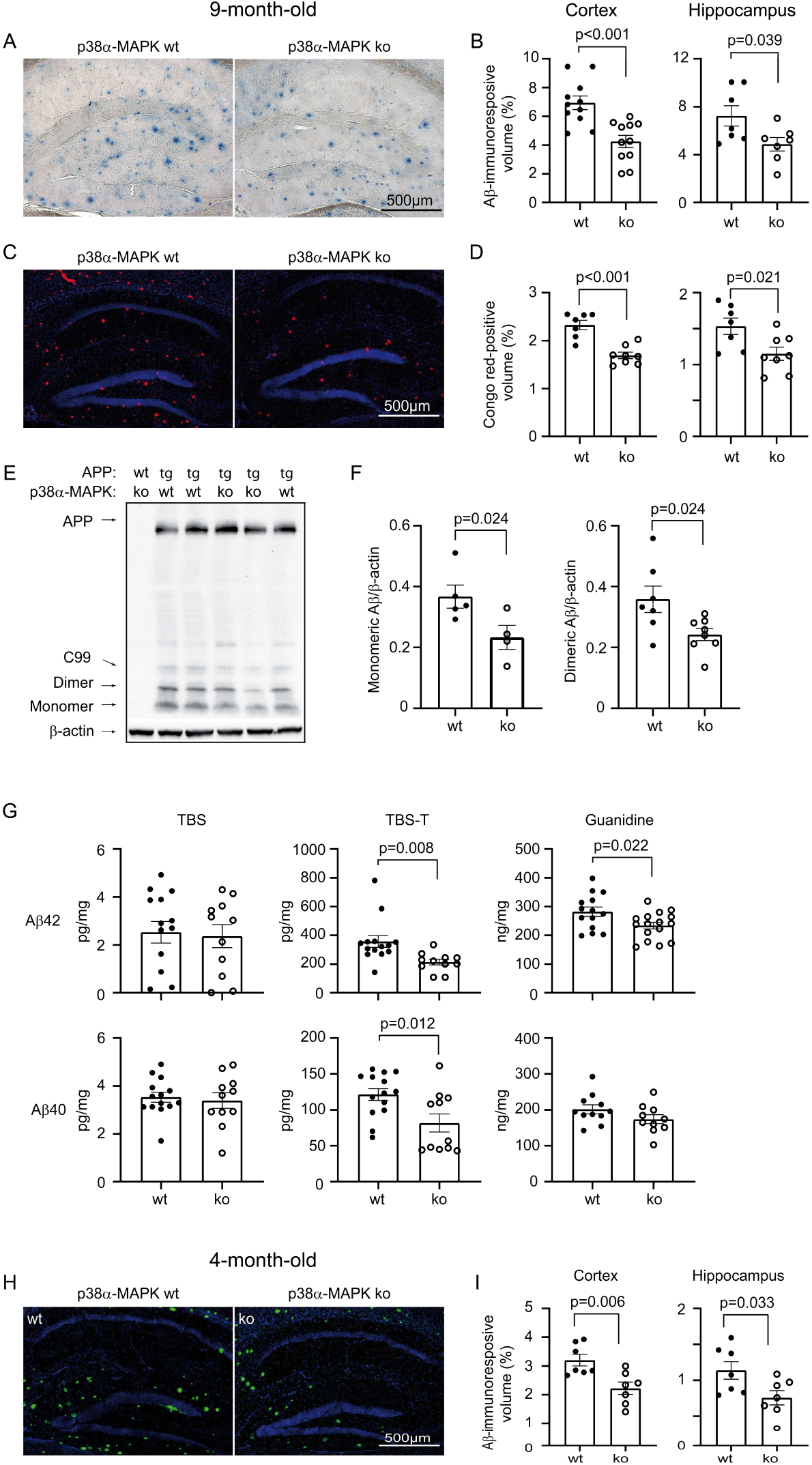
Deletion of p38α-MAPK in myeloid cells reduces cerebral Aβ load in APP-transgenic mice. Four and nine-month-old APP^tg^p38^fl/fl^LysMCre^+/-^ (p38α ko) and APP^tg^p38^fl/fl^LysMCre^-/-^ (p38α wt) were analyzed with stereological *Cavalieri* method for the cerebral Aβ volume (adjusted by the volume of analyzed tissues) after immunohistochemical (A and B), immunofluorescent (H and I) and Congo red (C and D) staining. Deletion of p38α-MAPK in myeloid cells significantly decreases Aβ deposits in the brain (B, D and I; *t* test; *n* ≥ 7 per each group). E and F, the brains derived from 9-month-old p38α-MAPK-ko and wt APP-transgenic mice were homogenized in RIPA buffer for Western blot analysis of soluble Aβ (monomeric and dimeric). Deficiency of p38α-MAPK significantly reduced soluble Aβ in the APP-transgenic mouse brain (*t* test; *n* ≥ 4 per each group). Moreover, APP-transgenic brains were homogenized and separated into TBS-, TBS-T-, and guanidine-soluble fractions. Amounts of Aβ40 and Aβ42 were measured by ELISA and normalized to the homogenate protein concentration. The concentrations of both Aβ40 and Aβ42 in TBS-T-soluble fractions and Aβ42 in guanidine-soluble fraction were significantly lower in myeloid p38α-ko APP mice than in p38α-wt control mice (G; *t* test; *n* ≥ 10 per group).

The amount of differently aggregated Aβ in brain tissue homogenates was measured with Western blot and ELISA. We observed that protein levels of monomeric and dimeric Aβ in 9-month-old APP^tg^p38^fl/fl^LysM-Cre^+/-^ mice were significantly lower than that in APP^tg^p38^fl/fl^LysM-Cre^-/-^ littermates (Fig. 2, E and F; *t* test, *p* < 0.05). Similarly, deletion of myeloid p38α-MAPK significantly decreased concentrations of both Aβ40 and Aβ42 in TBS plus 1% Triton X-100 (TBS-T)-soluble, and Aβ42 in guanidine hydrochloride-soluble brain homogenates from p38α-MAPK-deficient APP-transgenic mice compared with p38α-MAPK-wild-type littermate APP mice (Fig. 2, G; *t* test, *p* < 0.05).

In order to learn the effects of myeloid p38α-MAPK on Aβ pathology during the disease progression, we continued to analyze Aβ deposits in 4-month-old APP^tg^p38α^fl/fl^LysM-Cre^+/-^ and APP^tg^p38α^fl/fl^LysM-Cre^-/-^ mice. Deletion of p38α-MAPK in myeloid cells already reduced Aβ deposits in APP-transgenic mouse brain at this early disease stage as compared with p38α-MAPK-wild-type APP-transgenic mice (Fig. 2, H and I; *t* test, *p* < 0.05), although Western blot did not show difference in oligomeric Aβ levels (data not shown).

### Deletion of p38α-MAPK in myeloid cells inhibits inflammatory activation in the brain of APP-transgenic mice

Microglial inflammatory activation contributes to neuronal degeneration (Heneka et al., 2015). We observed that the number of Iba-1-immunoreactive cells (representing microglia/brain macrophages) in the whole hippocampus and cortex was significantly fewer in 9-month-old myeloid p38α-MAPK-deficient APP-transgenic mice than in p38α-MAPK-wild-type APP-transgenic littermates: APP^tg^p38α^fl/fl^LysM-Cre^+/-^ mice, 13.82 ± 0.65 × 10^3^ cells/mm^3^ in the hippocampus and 14.63 ± 0.56 × 10^3^ cells/mm^3^ in the cortex; APP^tg^p38α^fl/fl^LysM-Cre^-/-^ mice, 17.06 ± 0.81 × 10^3^ cells/mm^3^ in the hippocampus and 17.98 ± 0.96 × 10^3^ cells/mm^3^ in the cortex (Fig. 3, A and B; *t* test, *p* < 0.05). However, deficiency of p38α-MAPK in myeloid cells did not reduce Iba-1-positive cells in 9-month-old non-APP transgenic mice (Fig. 3, B; *t* test, *p* > 0.05). We also observed that the number of Iba-1-immunoreactive cells in the brain was significantly fewer in myeloid p38α-MAPK-deficient APP-transgenic mice than in p38α-MAPK-wild-type APP-transgenic littermates at 4 months of age (Fig. 3, C; *t* test, *p* < 0.05). To investigate the underlying mechanism which mediates the reduction of microglial number, we stained Iba-1 and Ki-67, a cell-proliferation marker, to identify proliferating microglia as we did in a previous study (Liu et al., 2013). We observed that deficiency of p38α-MAPK in myeloid cells significantly reduced Iba-1^+^Ki-67^+^ cells in both cortex and hippocampus of 4-but not 9-month-old APP-transgenic mice (Supplementary Fig. 3; *t* test, *p* < 0.05).

**Fig. 3.**
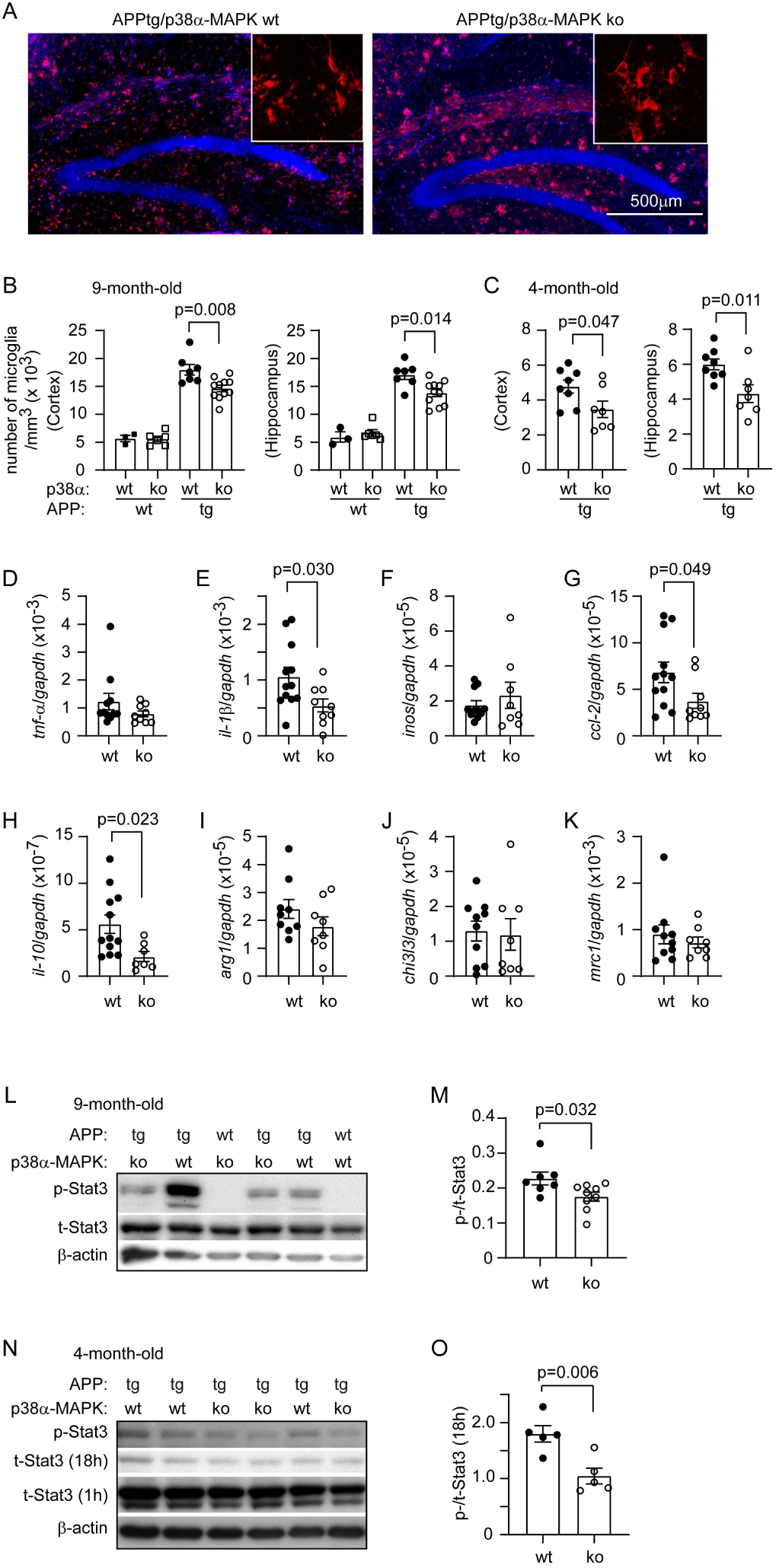
Deletion of p38α-MAPK in myeloid cells inhibits inflammatory activation in the brain of APP-transgenic mice. Four or nine-month-old APP-transgenic (APPtg) and non-transgenic (APPwt) mice with (p38α-ko) and without (p38α-wt) deletion of p38α-MAPK in the myeloid cell lineage were analyzed for the neuroinflammatory activation. Microglia were stained with fluorescence-conjugated Iba-1 antibodies (A, staining in 9-month-old APPtg mouse tissue) and counted with the stereological probe, Optical Fractionator. Deficiency of p38α-MAPK reduced Iba-1-positive cells in both 4 and 9-month-old APPtg mice, but not in 9-month-old APPwt mice (B; one-way ANOVA followed by Bonferroni *post hoc* test; *n* ≥ 7 per group for APPtg mice and ≥ 3 per group for APPwt mice. C; t test; *n* ≥ 7 per group). The inflammatory activation in brain was further analyzed with real-time PCR to detect transcripts of both pro- and anti-inflammatory genes. Transcription of *il-1β, ccl-2* and *il-10* genes was reduced by deletion of p38α-MAPK in 9-but not 4-month-old APPtg mice (D - K; t test; *n* ≥ 8 per group for 9-month-old APPtg mice). Four and nine-month-old APP^tg^p38^fl/fl^LysMCre^+/-^ and APP^tg^p38^fl/fl^LysMCre^-/-^ mice were further analyzed with quantitative Western blot for the levels of phosphorylated (Tyr705) (p-) and total (t-) Stat3 in the brain. Deficiency of myeloid p38α-MAPK significantly inhibits the activation of Stat3 (as shown by the ratio of p-/t-Stat3) in the brain of both 4 and 9-month-old APP-transgenic mice (L - O; t test; *n* = 5 and *n* ≥ 7 per group for 4- and 9-month-old mice, respectively). To prevent overexposure of film, the membrane for t-Stat3 blotting was washed for extra 18 hours (N).

We further measured transcripts of M1-inflammatory genes (*tumor necrosis factor-α* [*tnf-α*], *interleukin-1β* [*il-1β*], *inducible nitric oxide synthase* [*inos*], and *chemokine (C–C motif) ligand 2* [*ccl-2*]) and M2-inflammatory genes (*il-10, arginase 1, chitinase-like 3* [*chi3l3*], and *mannose receptor C type 1* [*mrc1*]) in brains of 4- and 9-month-old APP^tg^p38α^fl/fl^LysM-Cre^+/-^ and APP^tg^p38α^fl/fl^LysM-Cre^-/-^ mice. As shown in Fig. 3, E, G and H, cerebral *il-1β, ccl-2* and *il-10* transcripts were significantly reduced in 9-month-old APP^tg^p38α^fl/fl^Cre^+/-^ mice compared to their APP^tg^p38α^fl/fl^Cre^-/-^ littermate mice (*t* test, *p* < 0.05). However, deficiency of p38α-MAPK in myeloid cells did not change the transcription of all tested inflammatory genes in 4-month-old APP-transgenic mice (data not shown).

IL-10 activates signal transducers and activators of transcription 3 (Stat3) (Braun et al., 2013). As inhibition of IL-10-Stat3 signaling potentially promotes Aβ clearance in AD mice (Chakrabarty et al., 2015; Guillot-Sestier et al., 2015; Reichenbach et al., 2019), we examined Stat3 activation in the brain of AD animals. We observed that levels of phosphorylated Stat3 in both 4- and 9-month-old APP^tg^p38^fl/fl^LysM-Cre^+/-^ mice were significantly lower than that in APP^tg^p38^fl/fl^LysM-Cre^-/-^ littermate controls (Fig. 3, L - O; *t* test, *p* < 0.05). In the brain of 9-month-old non-APP-transgenic mice, phosphorylated Stat3 was undetectable (Fig. 3, L). Thus, IL-10/Stat3-mediated inflammatory signaling is activated in APP-transgenic mouse brain and is inhibited by p38α-MAPK deletion in myeloid cells in both early and late disease stages.

### Deletion of myeloid p38α-MAPK differently regulates microglial inflammatory activity in early and late disease stages of APP-transgenic mice

After observing that myeloid deficiency of p38α-MAPK attenuated both Aβ and inflammation in the whole brain, we asked how p38α-MAPK regulates inflammatory activation in individual microglial cells. We isolated CD11b^+^ microglia from both 4- and 9-month-old APP-transgenic mouse brains with magnetic beads-conjugated antibodies. Inflammatory gene transcripts were detected with quantitative RT-PCR. Surprisingly, deficiency of p38α-MAPK significantly reduced the transcription of *il-1β, ccl-2* and *il-10* genes in cells from 4-month-old APP-transgenic mice (Fig. 4, B, C and D; *t* test, *p* < 0.05), but increased the transcription of *il-1β* and *ccl-2* genes in cells from 9-month-old APP-transgenic mice (Fig. 4, F and G; *t* test, *p* < 0.05) as compared with p38α-MAPK-wildtype APP-transgenic mice.

**Fig. 4.**
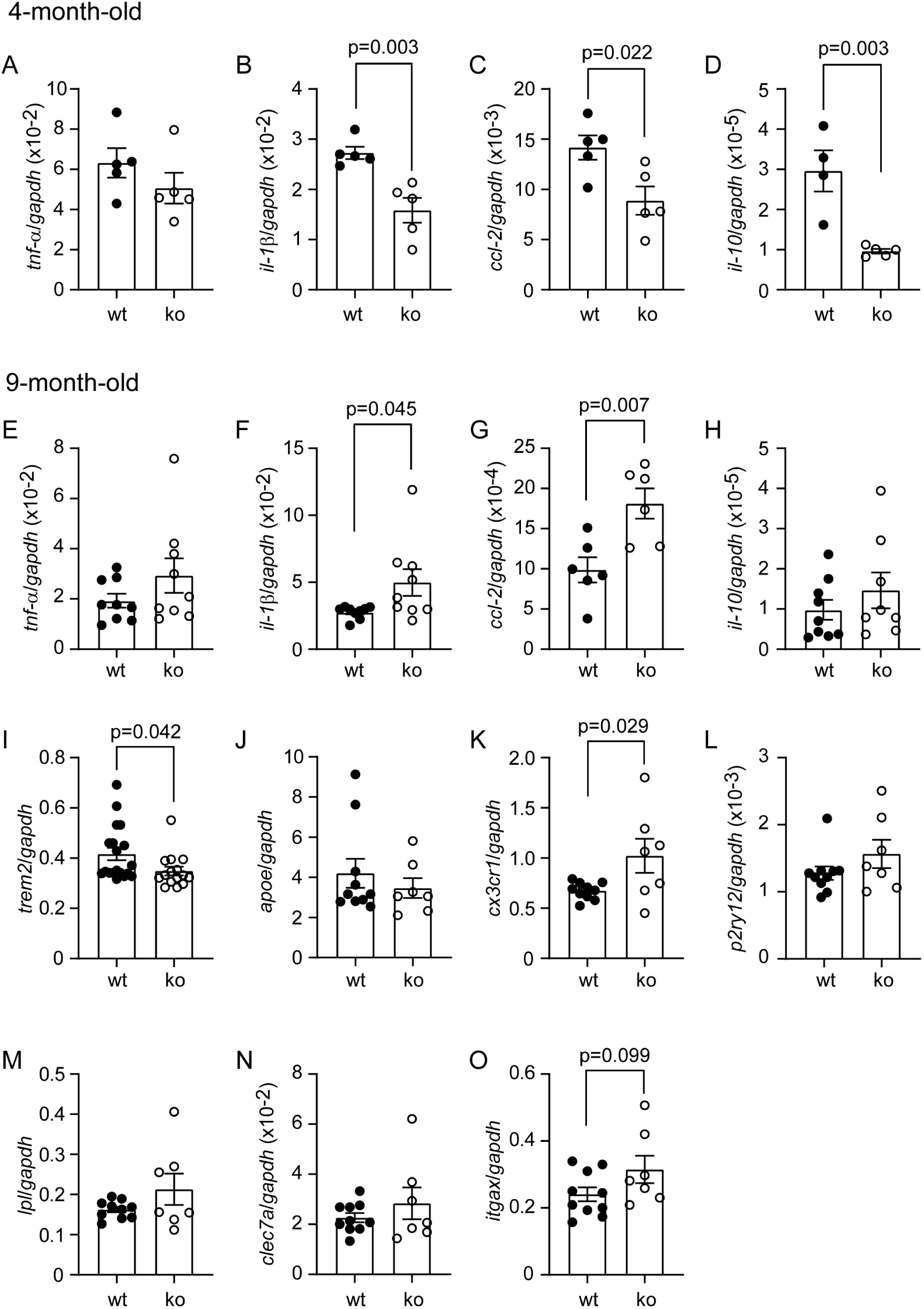
Deletion of myeloid p38α-MAPK differently regulates microglial inflammatory activity in the brain of APP-transgenic mice at early and late disease stages. Adult microglia were isolated with magnetic beads-conjugated CD11b antibodies from 4- and 9-month-old APP-transgenic mice with (p38α ko) and without (p38α wt) deletion of p38α-MAPK in myeloid cells. The transcripts of inflammatory genes and signature genes of disease-associated microglia in microglia were detected with real-time PCR. Deficiency of myeloid p38α-MAPK significantly inhibits transcription of inflammatory genes (e.g. *il-1β* and *ccl-2*) in individual microglia by 4 months of age but enhances it after the disease progresses (*t* test, *n* ≥ 4 per group for 4-month-old APP mice, and *n* ≥ 6 per group for 9-month-old APP mice).

Recently, disease-associated microglia (DAM) phenotype was defined after comparing microglial transcriptome between APP/PS1-transgenic and wild-type mice (Keren-Shaul et al., 2017; Krasemann et al., 2017). We detected transcripts of signature genes in isolated CD11b^+^ microglia from APP^tg^p38α^fl/fl^LysM-Cre^+/-^ and APP^tg^p38α^fl/fl^LysM-Cre^-/-^ littermate mice. We observed, in 9-month-old but not in 4-month-old APP-transgenic mice, that deletion of p38α-MAPK in myeloid cells: 1) significantly increased transcription of *cx3cr1* gene (Fig. 4, K; *p* = 0.029); 2) markedly deceased transcription of *trem2* gene (Fig. 4, I; *t* test, *p* = 0.042); and 3) tended to up-regulate the transcription of *itgax* gene (Fig. 4, O; *t* test, *p* = 0.099).

As disease progresses in APP^tg^p38^fl/fl^LysM-Cre^+/-^ mice, inflammatory stimuli accumulate, including aggregated Aβ and inflammatory molecules released either passively from injured/inflamed tissues or actively by activated microglia (e.g. IL-1β), although the inflammatory stimulation overall in APP^tg^p38^fl/fl^LysM-Cre^+/-^ mice might be weaker than in APP^tg^p38^fl/fl^LysM-Cre^-/-^ mice. We hypothesized that p38α-MAPK-mediated inflammatory regulation depends on the intensity of inflammatory stimulation. We cultured bone marrow-derived macrophages (BMDMs) from p38^fl/fl^LysM-Cre^+/-^ and p38^fl/fl^LysM-Cre^-/-^ mice. We observed that macrophages deleted for p38α-MAPK showed decreased TNF-α secretion, compared to p38α-MAPK-wildtype macrophages, when challenged with lipopolysaccharide (LPS) or Toll-like receptor 2 ligand, Pam3Cys-SKKKK, at low concentrations, but increased TNF-α secretion when challenged at higher concentrations (see supplementary Fig. 4, A and C). Deficiency of p38α-MAPK constantly decreased IL-10 secretion from macrophages treated with both LPS and Pam3Cys-SKKKK at various concentrations. Interestingly, the increase of TNF-α release was abolished by pre-treatment of recombinant IL-10 (see supplementary Fig. 4, D). This finding suggests that the inflammatory activity in p38α-MAPK-deficient microglia was shaped by the evolving extracellular inflammatory environment during the disease progresses.

### Deletion of myeloid p38α-MAPK potentially increases microglial clearance of Aβ in APP-transgenic mice at the late disease stage

Microglial uptake of Aβ is an important mechanism to clear Aβ in AD brain (Heneka et al., 2015). We asked whether deficiency of p38α-MAPK facilitates microglial internalization of Aβ in AD mice. After observing that there were more microglia surrounding Aβ deposits in 9-month-old APP^tg^p38^fl/fl^LysM-Cre^+/-^ mice than in APP^tg^p38^fl/fl^LysM-Cre^-/-^ littermate mice (Fig. 5, A and B; *t* test, *p* = 0.012), we isolated microglia from these two groups of AD mice with magnetic beads-conjugated CD11b antibodies and quantitate intracellular Aβ with Western blot. As shown in Fig. 5, C and D, the protein level of intracellular Aβ in p38α-MAPK-deficient cells was higher than that in p38α-MAPK-wild-type microglia (*t* test, *p* = 0.030). Furthermore, we evaluated expression levels of Aβ internalization-associated receptors in microglia with quantitative RT-PCR and flow cytometry. Deficiency of p38α-MAPK increased scavenger receptor A (SR-A) expression in microglia at both transcriptional and protein levels compared with microglia from p38α-MAPK-wildtype APP-transgenic mice (Fig. 5, E and I; *t* test, *p* < 0.05). The transcription of other Aβ internalization-associated receptors, such as CD36 and RAGE, was not significantly changed by deficiency of p38α-MAPK (Fig. 5, F and G; *t* test, *p* > 0.05).

**Fig. 5.**
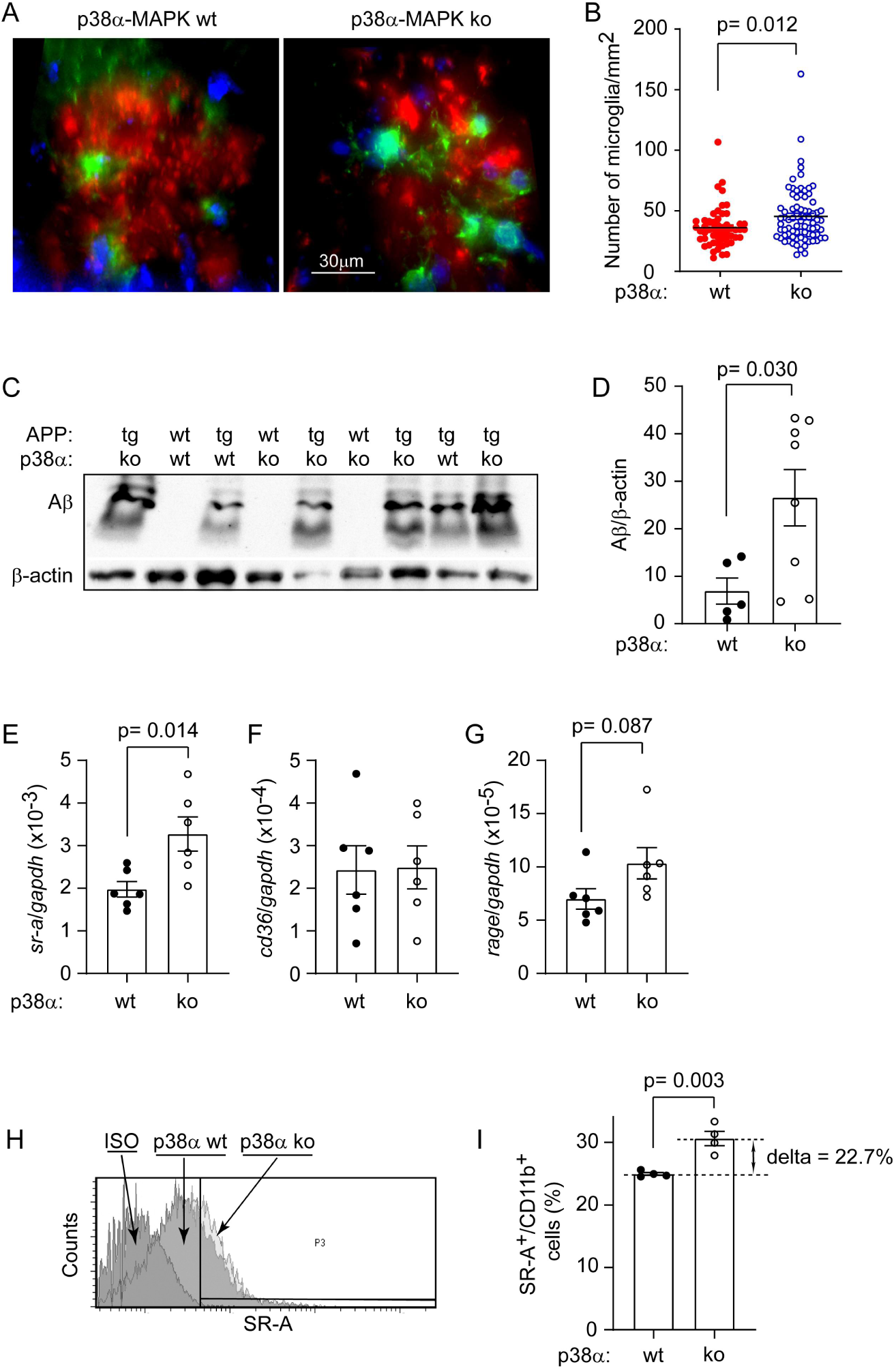
Deletion of p38α-MAPK in myeloid cells promotes microglial internalization of Aβ in the brain of 9-month-old APP-transgenic mice. Nine-month-old APP^tg^p38^fl/fl^LysMCre^+/-^ (p38α-MAPK ko, *n* = 5) and APP^tg^p38^fl/fl^LysMCre^-/-^ (p38α-MAPK wt, *n* = 4) mouse brains were stained for microglia with green fluorescence-conjugated Iba-1 antibodies and for Aβ deposits with Congo red (A). Under the red channel, total 105 Aβ deposits in p38α-MAPK ko mice and 76 Aβ deposits in p38α-MAPK wt mice were randomly chosen. The volume was estimated with *Cavalieri*’s method and the microglia with clear DAPI-stained nuclei and with contact to Aβ deposits were counted. Compared to p38α-MAPK wt mice, p38α-MAPK ko mice show significantly more microglia around Aβ deposits (B; *t* test). Adult microglia were also isolated with magnetic beads-conjugated CD11b antibodies from 9-month-old APP-transgenic mice with and without deletion of p38α-MAPK in myeloid cells. The intracellular Aβ of microglia was quantified using Western blot with human Aβ and β-actin antibodies. As a control, no Aβ was detected in the microglia isolated from APP-wild-type mice (C). The level of intracellular Aβ in p38α-MAPK ko cells is significantly higher than that in p38α-MAPK wt cells (D; *t* test, *n* = 5 for p38α-MAPK wt and 8 for p38α-MAPK ko microglia). The microglial gene transcription of Aβ internalization-associated receptors, such as SR-A, CD36, and RAGE, in 9-month-old APP-transgenic mice were detected with real-time PCR (E - G; *t* test, *n* ≥ 6 per group). Moreover, the protein level of SR-A on microglia was determined by flow cytometry after immunofluorescent staining of SR-A (H). At both transcriptional and protein levels, SR-A was up-regulated by deficiency of p38α-MAPK, as compared to p38α-MAPK-wild-type cells (E and I; *t* test, for flow cytometry, *n* = 4 per group).

In 4-month-old APP-transgenic mice, we repeated all experiments for 9-month-old mice. We observed that p38α-MAPK deficiency neither altered the intracellular Aβ in microglia, nor affected the transcription of Aβ internalization-associated receptors, including SR-A, CD36, and RAGE (data not shown).

In order to investigate whether inflammatory activation is essential for the enhancement of Aβ internalization in p38α-MAPK-deficient microglia, we cultured p38α-MAPK-deficient and wildtype BMDMs and primed them with and without 100 ng/ml LPS for 48 hours. Deficiency of p38α-MAPK did increase Aβ internalization in inflammatorily activated macrophages in association with an up-regulation of SR-A, but not in resting cells (see supplementary Fig. 5).

### Delayed deletion of p38α-MAPK specifically in microglia attenuates inflammation and Aβ deposits in the brain of aged APP-transgenic mice

Our study with APP^tg^p38^fl/fl^LysM-Cre^+/-^ mice and cultured macrophages indicated that p38α-MAPK deficiency enhanced inflammatory activity of microglia upon a strong inflammatory stimulation, which raised a question whether inhibition of p38α-MAPK in APP-transgenic mice starting at a late disease stage exaggerates the inflammatory activation in the whole brain, as substantial Aβ and microglia have been accumulated in the brain. It prevents p38α-MAPK inhibition from becoming an effective AD therapy. To answer this question, we constructed a second AD mouse model as we did in a recent study (Quan et al., 2021) by cross-breeding APP-transgenic mice with *mapk14*-floxed mice and Cx3Cr1-CreERT2 mice (Goldmann et al., 2013). We injected 9-month-old APP^tg^p38^fl/fl^Cx3Cr1-Cre^+/-^ and APP^tg^p38^fl/fl^Cx3Cr1-Cre^-/-^ littermate mice with tamoxifen, and analyzed mice by 12 months (Fig. 6, A). The efficiency of tamoxifen-induced gene recombination was 97% in CD11b+ microglia in APP^tg^p38^fl/fl^Cx3Cr1-Cre^+/-^ mice (*mapk14*/*gapdh*: 0.007 ± 0.001 and 0.212 ± 0.035, in APP^tg^p38^fl/fl^Cx3Cr1-Cre^+/-^ and APP^tg^p38^fl/fl^Cx3Cr1-Cre^-/-^ mice, respectively; *t* test, *p* < 0.001), which was in accordance with a previous observation (Goldmann et al., 2013). As a control, the transcriptional level of *mapk14* gene in CD11b+ blood cells was not different between these two groups of mice.

**Fig. 6.**
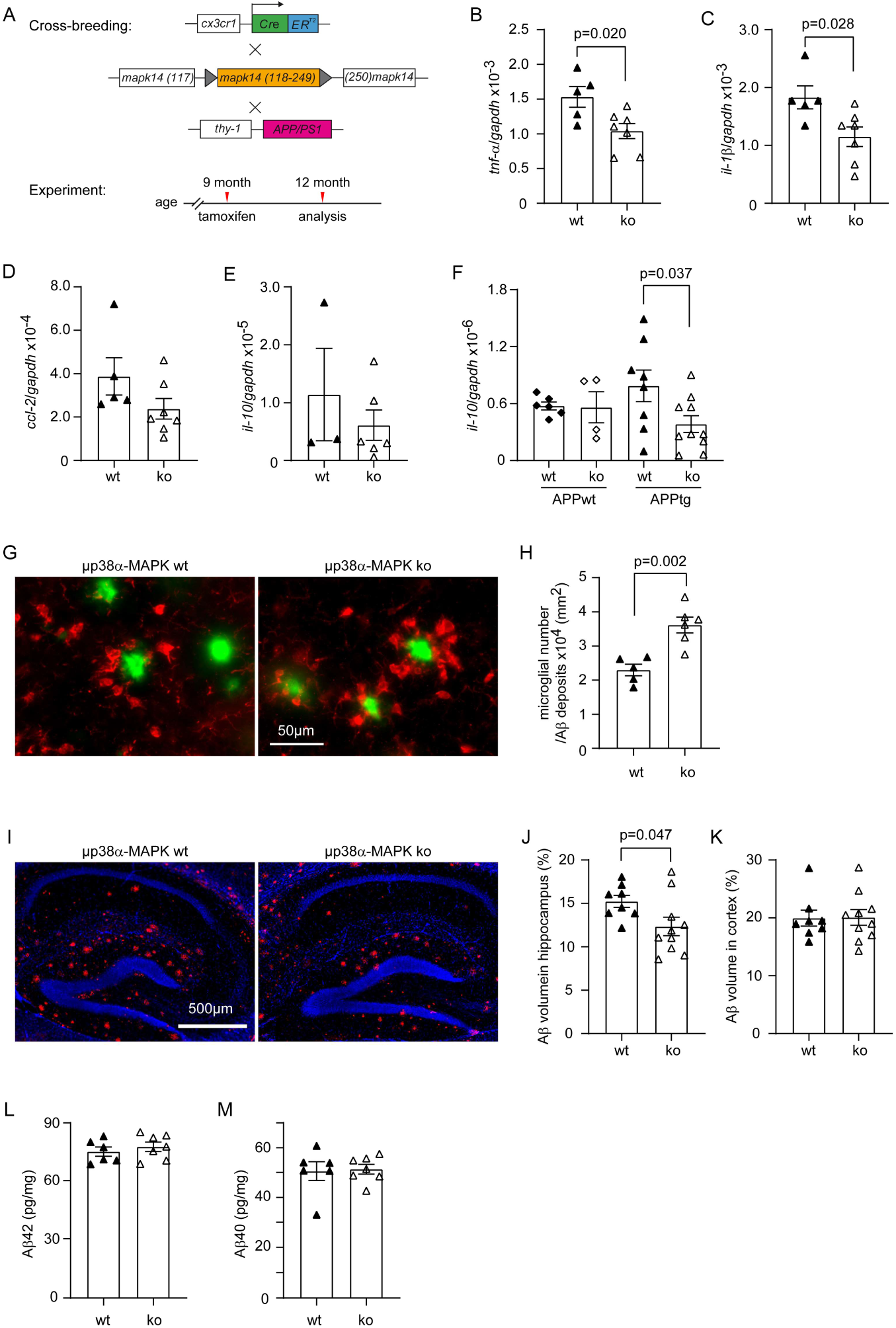
Deletion of p38α-MAPK specifically in microglia inhibits microglial inflammatory activation and decreases Aβ load in the brain of aged APP-transgenic mice. APP-transgenic mice (APPtg), *mapk14*-floxed mice and Cx3Cr1-CreERT2 mice were cross-bred. AD mice were injected with tamoxifen at 9 months of age for the induction of p38α-MAPK deletion in microglia and analyzed for AD pathology 3 months later (A). CD11b+ cells were isolated from brains of 12-month-old APPtg mice with (μp38α-ko) and without (μp38α-wt) deletion of p38α-MAPK in microglia, and detected with real-time PCR for the transcription of various inflammatory genes. Transcripts of *tnf-a* and *il-1β*, but not *ccl-2* and *il-10*, were significantly reduced in APPtg mice by microglial deficiency of p38α-MAPK (B and C; *t* test; *n* ≥ 5 per group). Deficiency of p38α-MAPK decreased the transcript of *il-10* in the brain tissue of 12-month-old APPtg mice compared with p38α-wt APPtg littermates (F; *t* test; *n* ≥ 8 per group). Twelve-month-old μp38α-wt and ko APPtg were counted for Iba-1-positive microglia (in red) around Aβ deposits stained with methoxy-XO4 (in green) (G). There are significantly more Aβ-associated microglia in the cortex and hippocampus of μp38α-ko APPtg mice than in μp38α-wt APP littermate controls (H; *t* test; *n* ≥ 5 per group). The coverage of Aβ deposits in the brain as stained by fluorescence-conjugated antibodies was further estimated with stereological *Cavalieri* method (I). Deletion of microglial p38α-MAPK reduces Aβ deposits in the hippocampus but not in the cortex of 12-month-old APP-transgenic mice (J and K; *t* test; *n* ≥ 8 per group). ELISA measurements of soluble Aβ40 and Aβ42 in RIPA-soluble brain homogenates did not show the difference of cerebral Aβ level between 12-month-old APP-transgenic mice with and without deletion of p38α-MAPK in microglia (L and M; *t* test, *p* > 0.05; *n* ≥ 6).

Surprisingly, deletion of p38α-MAPK in microglia of 12-month-old APP-transgenic mice (APP^tg^p38^fl/fl^Cx3Cr1-Cre^+/-^) compared with APP^tg^p38^fl/fl^Cx3Cr1-Cre^-/-^ littermates did not change the number of microglia in the hippocampus and cortex (data not shown); however, significantly decreased the transcripts of *tnf-α* and *il-1β*, but not *ccl-2, il-10, chi3l3* and *mrc1* genes in individual microglia (Fig. 6; B - E; *t* test, *p* < 0.05). In the brain tissue, deletion of p38α-MAPK did not change the transcription of inflammatory genes (e.g. *tnf-α, il-1β, ccl-2, chi3l3* and *mrc1*; data not shown), except that it decreased *il-10* gene transcription (Fig. 6; F; *t* test, *p* < 0.05). Deficiency of p38α-MAPK promoted Iba-1-positive microglia to cluster around Aβ deposits in both cortex and hippocampus as compared with p38α-MAPK-wild-type AD mice (Fig. 6, G and H; 3.61 ± 0.23 vs. 2.30 ± 0.17 ×10^4^ microglia/mm^2^ in p38α-MAPK-deficient and wildtype mice, respectively; *t* test, *p* = 0.002). We observed that deletion of p38α-MAPK by 9 months of age slightly but significantly decreased Aβ deposits in the hippocampus but not in the cortex (Fig. 6, I and J; *t* test, *p* = 0.047). Concentrations of both Aβ40 and Aβ42 in RIPA-soluble brain homogenates as quantitated by ELISA were not different between APP^tg^p38^fl/fl^Cx3Cr1-Cre^+/-^ and APP^tg^p38^fl/fl^Cx3Cr1-Cre^-/-^ mice (Fig. 6, L and M; *t* test, *p* > 0.05).

As APP^tg^p38^fl/fl^Cx3Cr1-Cre^+/-^ mice were haploinsufficient for *cx3cr1* gene, we performed additional experiments to examine whether the phenotype modification in APP^tg^p38^fl/fl^Cx3Cr1-Cre^+/-^ mice relative to APP^tg^p38^fl/fl^Cx3Cr1-Cre^-/-^ littermates was due to the haploinsufficiency of Cx3Cr1 rather than deficiency of p38α-MAPK in microglia. Our recent study demonstrated that Cx3Cr1 haploinsufficiency does not change Aβ load and neuroinflammation in APP-transgenic mice (Quan et al., 2021). In the current study, we showed that haploinsufficiency of Cx3Cr1 altered neither the recruitment of microglia toward Aβ deposits, nor the transcription of inflammatory genes, *tnf-α, il-1β, ccl-2* and *il-10* in individual microglia (see supplementary Fig. 6).

### Delayed deletion of p38α-MAPK in microglia has beneficial effects on neuronal protection in aged APP-transgenic mice

We further examined the effects of microglial deficiency of p38α-MAPK on neuronal function in aged APP-transgenic mice. We performed Morris water maze test to analyze the cognitive function of 12-month-old APP^tg^p38^fl/fl^Cx3Cr1-Cre^+/-^ and APP^tg^p38^fl/fl^Cx3Cr1-Cre^-/-^ mice and their APP-wildtype littermates. Deletion of microglial p38α-MAPK did not change the traveling distance and latency of APP-wildtype mice to reach the target platform (Fig. 7, A and B; two-away ANOVA, *p* > 0.05). However, APP^tg^p38^fl/fl^Cx3Cr1-Cre^+/-^ mice with deletion of p38α-MAPK specifically in microglia were able to find the hided platform with significantly less time and shorter traveling distance than APP^tg^p38^fl/fl^Cx3Cr1-Cre^-/-^ littermate mice (Fig. 7, A and B; two-away ANOVA, *p* < 0.05). In the 5-minute probe trial, APP^tg^p38^fl/fl^Cx3Cr1-Cre^+/-^ mice visited the region where the platform was previously located with significantly more frequency than APP^tg^p38^fl/fl^Cx3Cr1-Cre^-/-^ littermate mice (Fig. 7, D; one-away ANOVA, *p* < 0.05). Thus, deletion of microglial p38α-MAPK improved the cognitive function of aged AD mice.

**Fig. 7.**
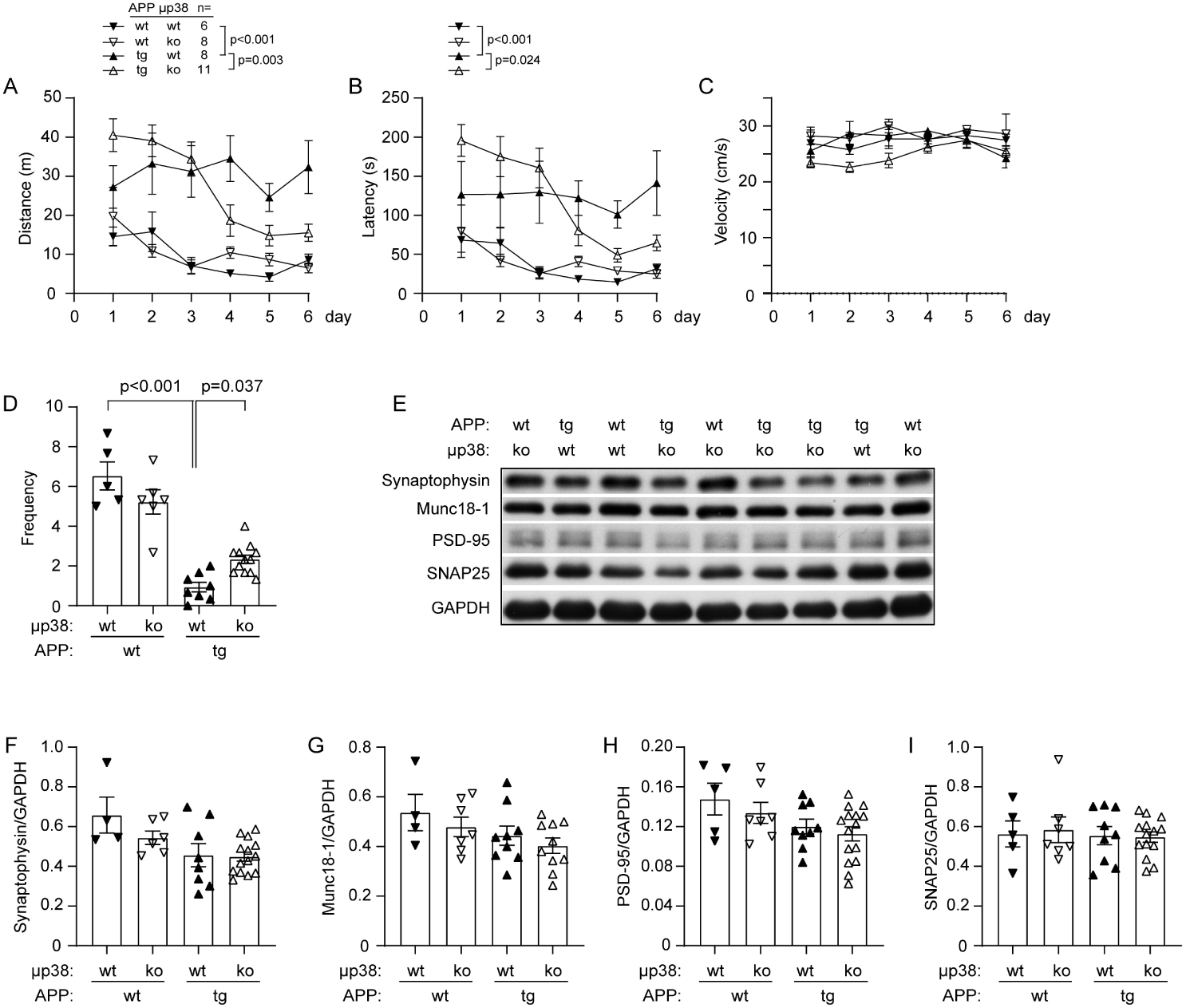
Deletion of p38α-MAPK in microglia improves cognitive function of aged APP-transgenic mice. During the training phase of water maze test, 9-month-old APP-transgenic mice (APPtg) spent significantly more time and traveled longer distances to reach the escape platform than did their non-APP-transgenic littermates (APPwt). Compared to mice with normal expression of p38α-MAPK (μp38α wt), deletion of p38α-MAPK in microglia (μp38α ko) significantly reduced the traveling time and distance of APPtg mice but not of APPwt mice after 3 days of training (A, B; two-way ANOVA from day 4 to day 6 followed by Bonferroni *post hoc* test; *n* is shown in the figure). Deficiency of microglial p38α-MAPK affected the swimming speed neither of APPtg and APPwt mice, nor for each mouse at different training time points (C; two-way ANOVA, *p* > 0.05). In the probe trial, APPtg mice crossed the platform region with significantly less frequency during the total 5-minute experiment; deletion of p38α-MAPK in microglia partially recovered APP-overexpression-induced cognitive impairments (D; one-way ANOVA followed by Bonferroni *post hoc* test). Western blotting was used to detect the amount of synaptic structure proteins, Munc18-1, SNAP25, synaptophysin, and PSD-95 in the brain homogenate of 12-month-old APPtg and APPwt mice (E - I). Deficiency in microglial p38α-MAPK changed the levels of all these 4 tested proteins neither in APPtg mice, nor in APPwt mice (one-way ANOVA followed by Bonferroni *post hoc* test; *n* ≥ 9 per group for APPtg mice and *n* ≥ 4 per group for APPwt mice).

However, when we detected synaptic proteins in the brain homogenates of 12-month-old APP^tg^p38^fl/fl^Cx3Cr1-Cre^+/-^ and APP^tg^p38^fl/fl^Cx3Cr1-Cre^-/-^ mice, we observed that microglial deficiency of p38α-MAPK by 9 - 12 months of age did not preserve the protein levels of synaptophysin, Munc18-1, PSD-95, and SNAP25 in APP-transgenic AD mice (Fig. 7, E - I; one-away ANOVA, *p* > 0.05).

### Deletion of p38α-MAPK in myeloid cells reduces IL17a-expressing lymphocytes and subsequently activates microglia and decreases Aβ in the brain of APP-transgenic mice

We observed different effects of p38α-MAPK deficiency in whole myeloid cells and specifically in microglia on microglial inflammatory activation and cerebral Aβ load. We supposed that p38α-MAPK-deficient peripheral myeloid cells play a distinct role. We isolated CD4+ spleen cells from 6-month-old APP-transgenic and wildtype mice and observed that transcription of *il-17a*, but not *ifn-γ, il-4* and *il-10* genes was significantly up-regulated in APP mice (see supplementary Fig. 7, A - D; *t* test, *p* < 0.05). By comparing gene transcripts in CD4+ spleen cells from 4- and 9-month-old APP^tg^p38^fl/fl^LysM-Cre^+/-^ and APP^tg^p38^fl/fl^LysM-Cre^-/-^ littermate mice, we observed that p38α-MAPK deficiency significantly reduced the transcription of *il-17a*, but not *ifn-γ, il-4* and *il-10* in AD mice at the age of 9, but not 4 months (Fig. 8, A - D; *t* test, *p* < 0.05).

**Fig. 8.**
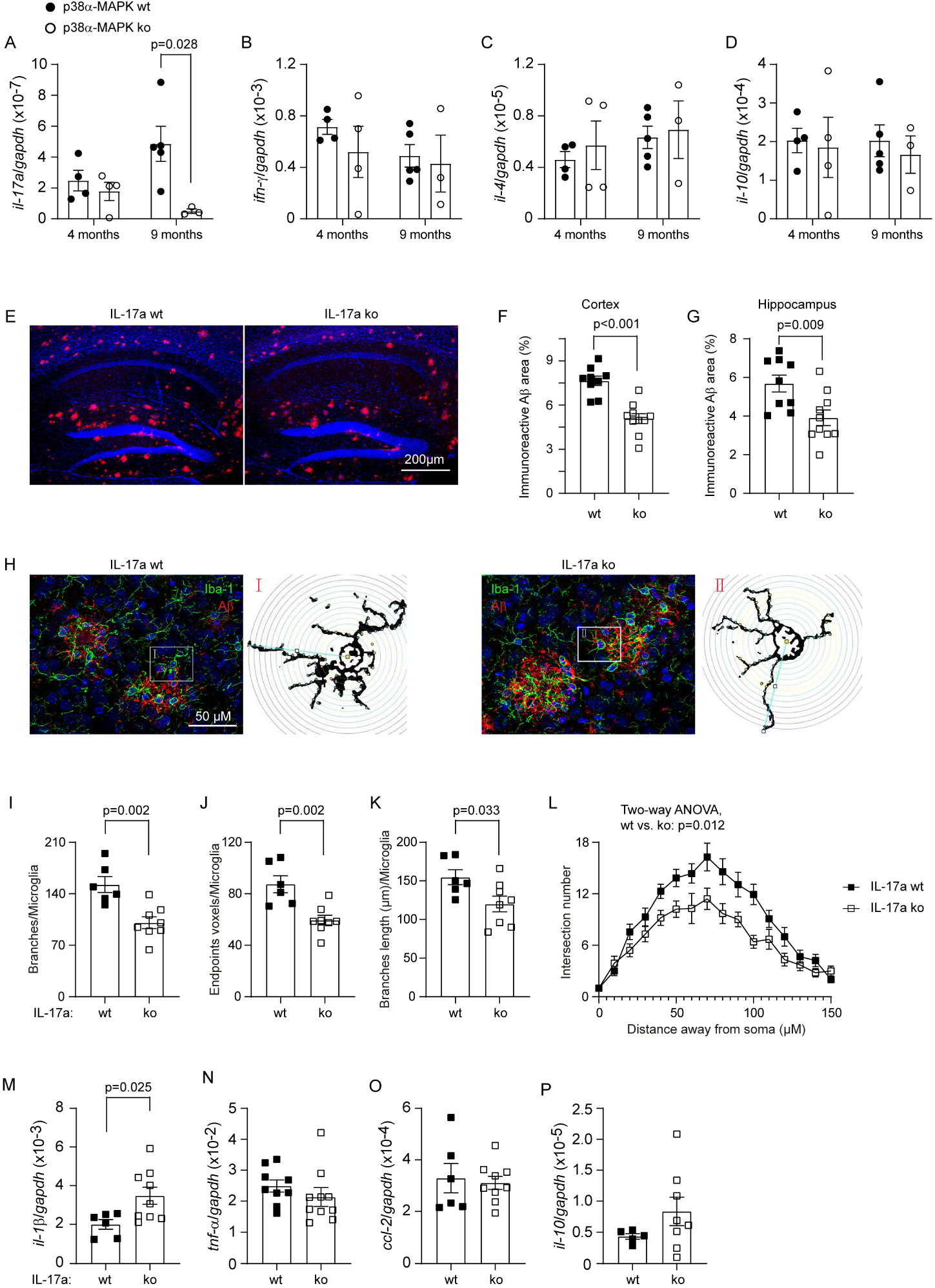
Deletion of p38α-MAPK in myeloid cells decreases *il-17a* gene transcription in CD4-positive spleen cells and IL-17a knockout activates microglia and reduces Aβ deposits in APP-transgenic mice. CD4-positive spleen cells were selected with magnetic beads-conjugated antibodies from 4- and 9-month-old APP-transgenic mice with (ko) and without (wt) deletion of p38α-MAPK in myeloid cells. Real-time PCR was used to quantify transcripts of marker genes for Th17 (*il-17a*), Th1 (*ifn-γ*), Th2 (*il-4*) and Treg lymphocytes (*il-10*) (A - D). Deletion of p38α-MAPK in myeloid cells down-regulates *il-17a* gene transcription in CD4-positive spleen cells from 9-but not 4-month-old APP-transgenic mice (A; *t* test between p38α-MAPK ko and wt mice, *n* ≥ 3 per group). To investigate the pathogenic role of IL-17a in AD, APP-transgenic mice were mated to IL-17a knockout mice. Brains of 6-month-old APP-transgenic mice with (ko) and without (wt) knockout of IL-17a were stained with antibodies against human Aβ and/or Iba-1 antibody (E and H). The area of immunoreactive Aβ-positive staining and the whole analyzed brain region was estimated with stereological *Cavalieri* method. IL-17a deficiency significantly decreases Aβ deposits in both cortex and hippocampus (F and G; *t* test, *n* ≥ 9 per group). The morphology of microglia in contact with Aβ deposits was analyzed (H). IL-17a deficiency significantly decreases the number and total length of branches of microglia (I - K; *t* test, *n* ≥ 6 per group; L; For the Sholl analysis, 10 microglia from 2 IL-17a ko mice and 14 microglia from 2 IL-17a wt mice were analyzed; two-way ANOVA was performed to test the difference between IL-17a ko and wt mice). CD11b-positive cells were further selected from 6-month-old IL-17a ko and wt APP-transgenic mice and detected for the inflammatory gene transcription with real-time PCR (M - P). IL-17a knockout significantly increases the transcripts of *il-1β* gene in CD11b-positive brain cells (M; *t* test, *n* ≥ 6 per group).

To investigate the role of IL-17a-expressing T lymphocytes in AD pathogenesis, we cross-bred APP-transgenic mice with *il-17a* knockout mice (Nakae et al., 2002). The coverage of immunoreactive Aβ deposits in both cortex and hippocampus of 6-month-old APP^tg^IL17a^-/-^(IL-17a knockout) mice was significantly less than in APP^tg^IL17a^+/+^ (IL-17a wildtype) littermates (Fig. 8, E - G; ko vs. wt: 5.08 % ± 0.34 % vs. 7.65 % ± 0.32 % in the cortex, and 3.91 % ± 0.40 % vs. 5.68 % ± 0.44 % in the hippocampus; *t* test, *p* < 0.001 and = 0.009, respectively).

By analyzing the morphology of microglia, we observed that IL-17a deficiency decreased the number, end points and length of branches of microglia, which had contacts with Aβ deposits (Fig. 8, H - K; *t* test, *p* < 0.001), but not of these microglia 50 μm away from the edge of Aβ plaques (data not shown). The Sholl analysis (Bird and Cuntz, 2019) showed that microglial branches crossing the concentric circles especially at around 70 μm from the soma was significantly fewer in APP^tg^IL17a^-/-^ than in APP^tg^IL17a^+/+^ mice (Fig. 8, L; two-away ANOVA, *p* < 0.05). Moreover, the transcriptional level of *il-1β* gene, but not *tnf-α, ccl-2, il-10* genes was higher in CD11b+ brain cells isolated from APP^tg^IL17a^-/-^ than that from APP^tg^IL17a^+/+^ mice (Fig. 8, M - P; *t* test, *p* < 0.05). Our results suggested that IL-17a deficiency had the potential to activate microglia and reduce Aβ load in the brain of AD mice, which mimicked the effects of p38α-MAPK-deficient myeloid cells in 9-month-old APP-transgenic mice.

## Discussion

In AD brain, microglial activation not only damages neurons by triggering neurotoxic inflammatory activation, but also saves neurons through clearing Aβ and p-tau (Heneka et al., 2015; Qin et al., 2016). We observed that deletion of p38α-MAPK in myeloid cell lineage or specifically in microglia decreases cerebral Aβ load, and improves the cognitive function of APP-transgenic mice. Deficiency of p38α-MAPK in myeloid cells inhibits inflammatory activation of individual microglia at an early disease stage, but enhances it after the disease progresses. Deletion of p38α-MAPK in peripheral myeloid cells potentially reduces IL17a-expressing lymphocytes, which subsequently activates microglia to clean Aβ in the brain.

We created two APP-transgenic mice with deletion of p38α-MAPK in the whole myeloid cells from birth (APP^tg^p38^fl/fl^LysM-Cre^+/-^) and specifically in microglia from 9 months of age (APP^tg^p38^fl/fl^Cx3Cr1-Cre^+/-^). Deficiency of p38α-MAPK in myeloid cells decreases cerebral Aβ load and protects neurons more efficiently than p38α-MAPK deficiency specifically in microglia. It is possible that an early deletion of p38α-MAPK in APP^tg^p38^fl/fl^LysM-Cre^+/-^ mice prevents Aβ deposition, and a late deletion of p38α-MAPK in APP^tg^p38^fl/fl^Cx3Cr1-Cre^+/-^ mice can only serve a limited effect on established Aβ plaques. However, it should be noted that p38α-MAPK-deficient peripheral myeloid cells might play an essential role in Aβ clearance in the brain. Peripheral myeloid cells have been discussed to regulate AD pathogenesis (Bettcher et al., 2021; Croese et al., 2021). In our AD model, there are around 2.5% peripheral myeloid cells potentially entering the brain, which are mainly located in proximity to blood vessels. The recruitment of myeloid cells is not strictly dependent on p38α-MAPK expression. Interestingly, we found that deficiency of p38α-MAPK in myeloid cells decreases the transcription of *il-17a* gene in CD4-positive spleen cells. As γδ T lymphocytes were not examined, we could not exclude that p38α-MAPK in myeloid cells also regulates the production of γδ T cells. We also observed that IL-17a knockout in APP-transgenic mice activates microglia and reduces Aβ in the brain. Thus, p38α-MAPK deficiency in peripheral myeloid cells potentially inhibits the generation of IL17a-expressing lymphocytes and modulates Aβ pathology in the brain. It has been reported that the number of Th17 cells increases in AD patients with MCI compared with non-dementia subjects or MCI patients due to other reasons (Oberstein et al., 2018). IL-17a-expressing T lymphocytes (mainly γδ T cells) accumulate in the meanings and brain of triple-transgenic AD mouse model (3xTg-AD). An intra-ventricle injection of IL-17-neutralizing antibody prevents the loss of cognitive function of mice (Brigas et al., 2021). The pathogenic role of IL-17a-expressing cells in AD should be further investigated in the following research.

Deficiency of p38α-MAPK in myeloid cells reduces neuroinflammation in APP-transgenic mice (APP^tg^p38^fl/fl^LysM-Cre^+/-^). Notably, p38α-MAPK deficiency in myeloid cells inhibits inflammatory activation in individual microglia at the early disease stage (by 4 months of age), but exaggerates microglial inflammatory activation after disease progression (by 9 month of age). The decrease of inflammatory gene transcripts in the whole brain of 9-month-old APP^tg^p38^fl/fl^LysM-Cre^+/-^ mice is likely due to the decreased number of microglia beginning earlier in the disease (e.g., The proliferation of microglia is reduced at 4 months of age), although the underlying mechanism is not yet understood. We also observed that p38α-MAPK deficiency inhibits pro-inflammatory activation in cultured macrophages challenged with TLR2 or TLR4 ligands at low concentrations, but enhances the inflammatory activation when challenged with higher concentrations of inflammatory stimuli. Thus, p38α-MAPK-mediated inflammatory regulation depends on the intensity of inflammatory stimulation. Our following experiments showed that: 1) p38α-MAPK deficiency in myeloid cells down-regulates *il-10* in the brain of AD mice; 2) p38α-MAPK deficiency reduces IL-10 secretion from LPS or Pam3Cys-SKKKK-treated cultured macrophages, and pre-treatment of IL-10 abolishes p38α-MAPK deficiency-enhanced TNF-α secretion from LPS-treated macrophages. It was reported that p38-MAPK drives expression of both pro- and anti-inflammatory cytokines, e.g. TNF-α, IL-1β and IL-10, in LPS-treated cultured macrophages; the released IL-10 thereafter dissolves the inflammatory activation by activating Stat3-Socs3 signaling pathway (Bode et al., 2012). Thus, an insufficiency of IL-10 in the brain of APP^tg^p38^fl/fl^LysM-Cre^+/-^ mice has a potential to promote the inflammatory activation of microglia. It is consistent with an observation that systemic stimulation with LPS increases TNF-α, IL-1β and IL-6 in the brain of myeloid p38α-MAPK-deficient mice in association with a decrease of IL-10 in the serum (Bachstetter et al., 2014). However, 12-month-old APP^tg^p38^fl/fl^Cx3Cr1-Cre^+/-^ mice with deletion of microglial p38α-MAPK from 9 months showed no increase in the inflammatory activation in microglia, although the neuroinflammatory activation in 12-month-old APP^tg^p38^fl/fl^Cx3Cr1-Cre^+/-^ mice should be stronger than in 9-month-old APP^tg^p38^fl/fl^LysM-Cre^+/-^ mice. It is unclear whether p38α-MAPK deficiency takes longer than 3 months to convert anti-inflammatory to pro-inflammatory activation in microglia.

The difference in microglial inflammatory activation between aged APP^tg^p38^fl/fl^LysM-Cre^+/-^ and APP^tg^p38^fl/fl^Cx3Cr1-Cre^+/-^ has yet to be clarified. It could again reinforce the role of IL-17a-expressing T lymphocytes in the regulation of microglia, since IL-17a knock-out increases the inflammatory activity of microglia in AD mice. The inflammatory enhancement of microglia is correlated with the reduced transcription of *il-17a* in CD4-positive spleen in 9-month-old APP^tg^p38^fl/fl^LysM-Cre^+/-^ mice. By 4 months, the *il-17a* transcription is not changed in spleen cells and the inflammatory activation of microglia is inhibited in APP^tg^p38^fl/fl^LysM-Cre^+/-^ mice.

The inflammatory activation might promote Aβ clearance by p38α-MAPK-deficient microglia in AD brain. Microglia internalize more Aβ in APP^tg^p38^fl/fl^LysM-Cre^+/-^ mice than in APP^tg^p38^fl/fl^LysM-Cre^-/-^ littermates by 9 but not 4 months of age, which is associated with an increase of inflammatory activation. LPS priming is essential for p38α-MAPK deficiency to facilitate the uptake of Aβ by cultured macrophages. The upregulation of the expression of SR-A, a typical Aβ-phagocytic receptor (Paresce et al., 1996), could partially mediate the effect of p38α-MAPK deficiency. However, the inflammatory regulation of Aβ clearance is inconclusive. Pro-inflammatory activation inhibits Aβ uptake by cultured microglia and macrophages (Hao et al., 2011; Koenigsknecht-Talboo and Landreth, 2005; Liu et al., 2014); while, transgenic expression of TNF-α, IL-1β or IL-6 decreases Aβ in the brain of AD mice (Chakrabarty et al., 2011; Chakrabarty et al., 2010; Shaftel et al., 2007). Administration of TREM2 antibodies decreases Aβ load also with increased expression of inflammatory cytokines and chemokines in APP-transgenic mouse brain (Price et al., 2020; Wang et al., 2020).

In both APP^tg^p38^fl/fl^LysM-Cre^+/-^ and APP^tg^p38^fl/fl^Cx3Cr1-Cre^+/-^ mice, p38α-MAPK deficiency promotes microglial recruitment around Aβ deposits, which favors Aβ clearance (Hao et al., 2011; Liu et al., 2014; Quan et al., 2021). Microglia clustering around Aβ deposits was also considered to protect local neurite from damage by forming a physical barrier and compacting Aβ into dense plaques (Condello et al., 2015; Spangenberg et al., 2019; Wang et al., 2016; Yuan et al., 2016). However, the mechanism that drives microglia to migrate to Aβ deposits is unclear. We observed that IL-10-Stat3 signaling is inhibited in the brain of p38α-MAPK-deficient APP-transgenic mice. It has been observed that a lack of IL-10 or Stat3 facilitates microglial recruitment to Aβ deposits and increases Aβ clearance in AD mice (Chakrabarty et al., 2015; Guillot-Sestier et al., 2015; Reichenbach et al., 2019). It again suggests that IL-10 might mediate the effect of p38α-MAPK in AD pathogenesis.

LysM-Cre was reported to mediate the gene recombination in neurons (Orthgiess et al., 2016). However, many other studies showed very few or no neurons exhibiting LysM promoter activity (Goulielmaki et al., 2020; Liu et al., 2019). We observed that LysM-Cre drives expression of the reporter, GFP, in few neurons in the hippocampus, but not in neurons of the cortex, which suggests that LysM promoter is active only in a small subgroup of neurons. The change in the phenotype of APP^tg^p38^fl/fl^LysM-Cre^+/-^ mice compared to APP^tg^p38^fl/fl^LysM-Cre^-/-^ littermates results mainly from the lack of p38α-MAPK in myeloid cells. The enhancement of microglial inflammatory activation in 9-month-old APP^tg^p38^fl/fl^LysM-Cre^+/-^ mice does not support a substantial deletion of p38α-MAPK in neurons, since a deficiency of p38α-MAPK in neurons reduces Aβ generation, which in principle inhibits the inflammatory activation of microglia (Schnöder et al., 2020; Schnöder et al., 2016).

Until now, clinical interventions to decrease inflammatory activation or reduce Aβ accumulation or p-tau aggregation in the brain (Doody et al., 2014; Egan et al., 2018; Gauthier et al., 2016; Group, 2015; Group et al., 2008) have not led to AD therapies efficiently modifying the disease progression. It is important to recognize that AD is a heterogeneous disease. We have observed that neuronal deficiency of p38α-MAPK reduces Aβ deposition and p-tau aggregation in both APP- and tau-transgenic mouse brains, which leads to improved cognitive function in AD mice (Schnöder et al., 2020; Schnöder et al., 2016; Schnöder et al., 2021). Our current study further shows that a p38α-MAPK deficiency in myeloid cells attenuates cerebral Aβ and cognitive deficits in APP-transgenic mice. We believe that p38α-MAPK is a novel target for AD therapy, by which all Aβ, p-tau and neuroinflammation-mediated AD pathogenic processes can be appropriately modified at the same time.

In summary, p38α-MAPK deficiency in peripheral myeloid cells instead of in microglia triggers an efficient Aβ clearance in the brain and improves cognitive function of APP-transgenic mice. Peripheral p38α-MAPK-deficient myeloid cells have a potential to decrease the generation of IL17a-expressing T lymphocytes, which subsequently activates microglia to internalize Aβ. To investigate the pathogenic mechanisms of IL-17a-expressing T lymphocytes and optimize AD therapy with p38α-MAPK inhibitors will be the focus of our following studies.

## Materials and methods

### Animal models and cross-breeding

APP/PS1-double transgenic mice (APP) over-expressing human mutated APP (KM670/671NL) and PS1 (L166P) under Thy-1 promoters (Radde et al., 2006) were kindly provided by M. Jucker, Hertie Institute for Clinical Brain Research, Tübingen, Germany; p38^fl/fl^ mice with loxP site-flanked *mapk14* gene (Nishida et al., 2004) were imported from BioResource Center, RIKEN Tsukuba Institute, Japan; Cx3Cr1-CreERT2 mice that express a fusion protein of Cre recombinase and an estrogen receptor ligand binding domain under the control of endogenous *cx3cr1* promoter/enhancer elements (Goldmann et al., 2013) were kindly provided by M. Prinz, University of Freiburg, Germany; and LysM-Cre knock-in mice expressing Cre from the endogenous lysozyme 2 gene locus (Clausen et al., 1999) were bought from The Jackson Laboratory, Bar Harbor, ME (stock number 004781) and were back-crossed to C57BL/6J mice for > 6 generations. APP-transgenic mice deficient of p38α-MAPK specifically in myeloid cells (e.g., microglia, macrophages and neutrophils) (APP^tg^p38^fl/fl^LysM-Cre^+/−^) were established by cross-breeding APP-transgenic mice with p38^fl/fl^ and LysM-Cre mice. To create AD mice with p38α-MAPK deficiency specifically in endogenous microglia (APP^tg^p38^fl/fl^Cx3Cr1-Cre^+/−^), APP-transgenic mice were first cross-bred with p38^fl/fl^ and Cx3Cr1-Cre mice and induced for the deletion of *mapk14* gene by intraperitoneal injection of tamoxifen (Sigma-Aldrich; 100 mg/kg) in corn oil once a day over 5 days. To delete IL-17a in AD mice, APP-transgenic mice were cross-bred with *il-17a* knockout mice (Nakae et al., 2002), which were kindly provided by Y. Iwakura, Tokyo University of Science, Japan. All mice from the same litter were used for experiments without any exclusion so that the phenotype of APP-transgenic mice with or without p38α-MAPK ablation was compared only between siblings. All animal experiments were performed in accordance with relevant national rules and authorized by Landesamt für Verbraucherschutz, Saarland, Germany (registration numbers: 40/2014, 12/2018 and 34/2019).

### Morris water maze

The Morris water maze test, consisting of a 6-day training phase and a 1-day probe trial, was used to assess the cognitive function of APP^tg^ mice and their wild-type littermates with and without deficiency of p38α-MAPK, as previously described (Qin et al., 2016). During training phase, latency time, distance, and velocity were recorded with Ethovision video tracking equipment and software (Noldus Information Technology, Wageningen, the Netherlands). During the probe trial, the platform was removed and we measured the latency of first visit to the location of original platform, the frequency of crossing in that location, and the time spent in the platform area.

### Collection of brain tissue

Animals were euthanized at the end experiments by inhalation of isoflurane. Mice were then perfused with ice-cold PBS, and the brain was removed and divided. The left hemisphere was immediately fixed in 4% paraformaldehyde (PFA; Sigma-Aldrich Chemie GmbH, Taufkirchen, Germany) for immunohistochemistry. A 0.5-μm-thick piece of tissue was sagittally cut from the right hemisphere. The cortex and hippocampus were carefully separated and homogenized in TRIzol (Thermo Fisher Scientific, Darmstadt, Germany) for RNA isolation. The remainder of the right hemisphere was snap-frozen in liquid nitrogen and stored at −80°C until biochemical analysis.

### Positive selection of CD11b or CD4-positive cells

To determine the gene expression and cell surface antigen expression in microglia/brain macrophages, we carefully dissected the entire cerebrum from 4- or 9-month-old APP-transgenic mice with or without a deficiency of p38α-MAPK and prepared a single-cell suspension. As described in a previous study (Liu et al., 2014), CD11b+ cells were selected with MicroBeads-conjugated CD11b antibody (clone M1/70.15.11.5; Miltenyi Biotec B.V. & Co. KG, Bergisch Gladbach, Germany). Lysis buffer was immediately added to CD11b+ cells for isolation of total RNA with the RNeasy Plus Mini Kit (Qiagen, Hilden, Germany); alternatively, CD11b+ cells were used for flow cytometric analysis after cells were stained with Alexa647-conjugated rat anti-mouse CD204 antibody (clone 2F8; Bio-Rad Laboratories GmbH, Feldkirchen, Germany) or CD11b+ cells were used for Western blot detection of Aβ after cells were lysed in radioimmunoprecipitation assay buffer (RIPA buffer; 50mM Tris [pH 8.0], 150mM NaCl, 0.1% SDS, 0.5% sodiumdeoxy-cholate, 1% NP-40, and 5mM EDTA) supplemented with protease inhibitor cocktail (Roche Applied Science, Mannheim, Germany). CD11b+ cells were also selected with magnetic beads-conjugated antibodies from the anticoagulated blood of AD mice.

To select CD4+ lymphocytes, we prepared single-cell suspensions from spleens of 4- and 12-month-old APP^tg^p38^fl/fl^LysM-Cre^+/-^ and APP^tg^p38^fl/fl^LysM-Cre^-/-^ littermate mice with the Spleen Dissociation Kit (Miltenyi Biotec B.V. & Co. KG). After blocking with 25 μg/ml rat anti-mouse CD16/CD32 antibody (clone 2.4G2; BD Biosciences, Heidelberg, Germany), we selected cells with Dynabeads®-conjugated antibody against mouse CD4 (L3T4) (Thermo Fisher Scientific). Lysis buffer was immediately added to CD4+ cells for isolation of RNA.

### Histological analysis

PFA-fixed left hemisphere was embedded in paraffin and serial 40-μm-thick sagittal sections were cut and mounted on glass slides. For each animal, 4 sections with an interval of 10 layers between neighboring sections were examined. Human Aβ was stained with rabbit anti-human Aβ antibody (clone D12B2; Cell Signaling Technology Europe, Frankfurt am Main, Germany) and microglia labeled with rabbit anti-Iba-1 antibody (Wako Chemicals, Neuss, Germany), and visualized with the VectaStain ABC-AP kit and the VECTOR Blue Alkaline Phosphatase Substrate kit (both from Vector Laboratories, Burlingame, USA) or fluorescence-conjugated second antibodies. Compacted Aβ in the brain tissue was stained with Congo red (Sigma-Aldrich Chemie GmbH) according to our established protocol (Liu et al., 2014). In the whole hippocampus and cortex, volumes of Aβ were estimated with the *Cavalieri* method, and Iba-1-positive cells were counted with Optical Fractionator as described previously (Liu et al., 2014) on a Zeiss AxioImager.Z2 microscope (Carl Zeiss Microscopy GmbH, Göttingen, Germany) equipped with a Stereo Investigator system (MBF Bioscience, Williston, USA).

The relationship between microglia and Aβ deposits was investigated as we did in a previous study (Hao et al., 2011). Serial brain sections were co-stained with Iba-1 antibody, and methoxy-XO4 (Bio-Techne GmbH, Wiesbaden, Germany) or Congo red. Under Zeiss microscopy with 40× objective, Aβ deposits and surrounding microglia were imaged with Z-stack serial scanning from −10 to +10 μm. From each section, ≥ 10 randomly chosen areas were analyzed. The total number (> 200) of Iba-1-positive cells co-localizing with Aβ deposits were counted. The area of Aβ was measured with Image J (https://imagej.nih.gov/ij/) for the adjudgment of microglial cell number.

### Analysis of microglial morphology

For the analysis of microglial morphology, published protocols and Fiji Image J were used (Fernandez-Arjona et al., 2017; Young and Morrison, 2018). PFA-fixed brain tissues were embedded in Tissue-Tek® O.C.T. Compound (Sakura Finetek Europe B.V., AJ Alphen aan den Rijn, the Netherlands). The brain was cut in 30-μm sagittal sections. After fluorescent staining with Iba-1 antibody and methoxy-XO4 (or Aβ antibody), total 10 Aβ plaques/mouse were randomly selected from the cortex dorsal to hippocampus and imaged under 40× objective with Z-stack scanning with 1 μm of interval. The serial images were Z-projected with maximal intensity, 8-bit grayscale transformed, Unsharp-Mask filter and despeckle-treated, and binarized to obtain a black and white image. The cells with complete nucleus and branches and without overlapping with neighboring cells were chosen for analysis. The single-pixel background noise was eliminated and the gaps along processes were filled under the view of the original image of the cell. The processed image was skeletonized and analyzed with the plugin Analyze Skeleton (2D/3D) (http://imagej.net/AnalyzeSkeleton) for the total number of primary branches, length of all branches, and the number of branch endpoints of each microglia. The whole analysis was done blinded to genotypes.

The Sholl analysis was performed using the Fiji Image J plugin Simple Neurite Tracer (https://imagej.net/Simple_Neurite_Tracer) and published protocols (Longair et al., 2011; Tavares et al., 2017), by counting the number of microglial branches at a particular distance from the soma. Total 10 microglia from 2 APP^tg^IL17a^-/-^ mice and 14 microglia from 2 APP^tg^IL17a^+/+^ littermates were analyzed. Concentric circles (radius) were set with 10 μm of interval from the soma to the longest branch.

### Western blot analysis

Frozen brain tissues were homogenized on ice in RIPA buffer supplemented with protease inhibitor cocktail (Roche Applied Science) and phosphatase inhibitors (50nM okadaic acid, 5mM sodium pyrophosphate, and 50mM NaF; Sigma-Aldrich). The protein levels of synaptic proteins: Munc18-1, synaptophysin, SNAP-25, and PSD-95, and β-actin or α-tubulin as a loading control, were detected with quantitative Western blot as we did in previous studies (Quan et al., 2021; Schnöder et al., 2020). For quantification of phosphorylated and total amount of Stat3, brain homogenates were prepared from 4- and 9-month-old p38α-MAPK deficient and wildtype AD mice and detected with rabbit or mouse monoclonal antibodies against phosphorylated and total Stat3, and β-actin (clone D3A7, 124H6 and 13E5, respectively; all antibodies were bought from Cell Signaling Technology).

### Brain homogenates and Aβ ELISA and Western blot analysis

The frozen brain hemispheres were homogenized and extracted serially in TBS, TBS plus 1% Triton X-100 (TBS-T), guanidine buffer (5 M guanidine HCl/50 mM Tris, pH 8.0) as we did in a previous study (Liu et al., 2014). Aβ concentrations in three separate fractions of brain homogenates were determined by Aβ42 and Aβ40 ELISA kits (both from Thermo Fisher Scientific). Results were normalized on the basis of the sample’s protein concentration.

For detection of Aβ oligomers with our established method (Schnöder et al., 2016), the brain was homogenized in RIPA buffer. The protein was separated by 10 - 20% pre-casted Tris-Tricine gels (Anamed Elektrophorese GmbH, Groß-Bieberau/Rodau, Germany). Anti-human Aβ mouse monoclonal antibody (clone W0-2; Merck Chemicals GmbH, Darmstadt, Germany) and rabbit anti-β-actin monoclonal antibody (clone 13E5; Cell Signaling Technology) were used for Western blot.

### Quantitative PCR for analysis of gene transcripts

Total RNA was isolated from mouse brains with TRIzol or from selected cells with RNeasy Plus Mini Kit (Qiagen) and reverse-transcribed. Gene transcripts were quantified with established protocols (Liu et al., 2014; Quan et al., 2021) and Taqman gene expression assays of mouse *tnf*-*α, il-1β, inos, ccl-2, il-10, arginase 1, chi3l3, mrc1, apoe, trem2, p2ry12, cx3cr1, lpl, clec7a, itgax, sr-a, cd36, rage* and *gapdh* (Thermo Fisher Scientific). The transcription of *mapk14* genes in CD11b-positive cells was determined using the SYBR green binding technique with the following primers: 5’-CCCGAACGATACCAGAACCT-3’ and 5’-CTTCAGCAGACGCAACTCTC-3’.

### Statistical analysis

Data were presented as mean ± SEM. For multiple comparisons, we used one-way or two-way ANOVA followed by Bonferroni, Tukey, or Dunnett T3 *post hoc* test (dependent on the result of Levene’s test to determine the equality of variances). Two independent-samples Students *t*-test was used to compare means for two groups of cases. All statistical analyses were performed with GraphPad Prism 8 version 8.0.2. for Windows (GraphPad Software, San Diego, USA). Statistical significance was set at p < 0.05.

## Acknowledgments

We thank Dr. M. Jucker (Hertie Institute for Clinical Brain Research, Tübingen) for providing APP/PS1-transgenic mice, Dr. M. Prinz (Department of Neuropathology, University of Freiburg, Freiburg) for Cx3Cr1-CreERT2 mice, and S. Offermanns (Max Planck Institute for Heart and Lung Research) for *gpr43*-floxed mice. The floxed-p38α-MAPK mice were kindly provided by Dr K. Otsu (Osaka University) through the RIKEN Bioresource Center. We appreciate Elisabeth Gluding and Isabel Euler for their excellent technical assistance. This work was supported by Deutsche Forschungsgemeinschaft (LI1725/2-1; to Y.L.); Alzheimer Forschung Initiative e.V. (#18009; to Y.L.) and Saarland University through Anschubfinanzierung 2021 (to Y.L.).

## Author Contributions

Y.L. conceptualized and designed the study, acquired funding, conducted experiments, acquired and analyzed data, and wrote the manuscript. Q.L., L.S., W.H., K.L., Y.D., and I.T. conducted experiments, acquired data and analyzed data. M.M. offered an animal facility and supervised animal experiments. K.F. offered a research laboratory and supervised the laboratory work. All authors contributed to the article and approved the submitted version.

## Data Availability Statement

The data that support the findings of this study are available from the corresponding author upon reasonable request.

