## Supplementary figures for "p38α-MAPK-deficient myeloid cells ameliorate symptoms and pathology of APP-transgenic AD mice"

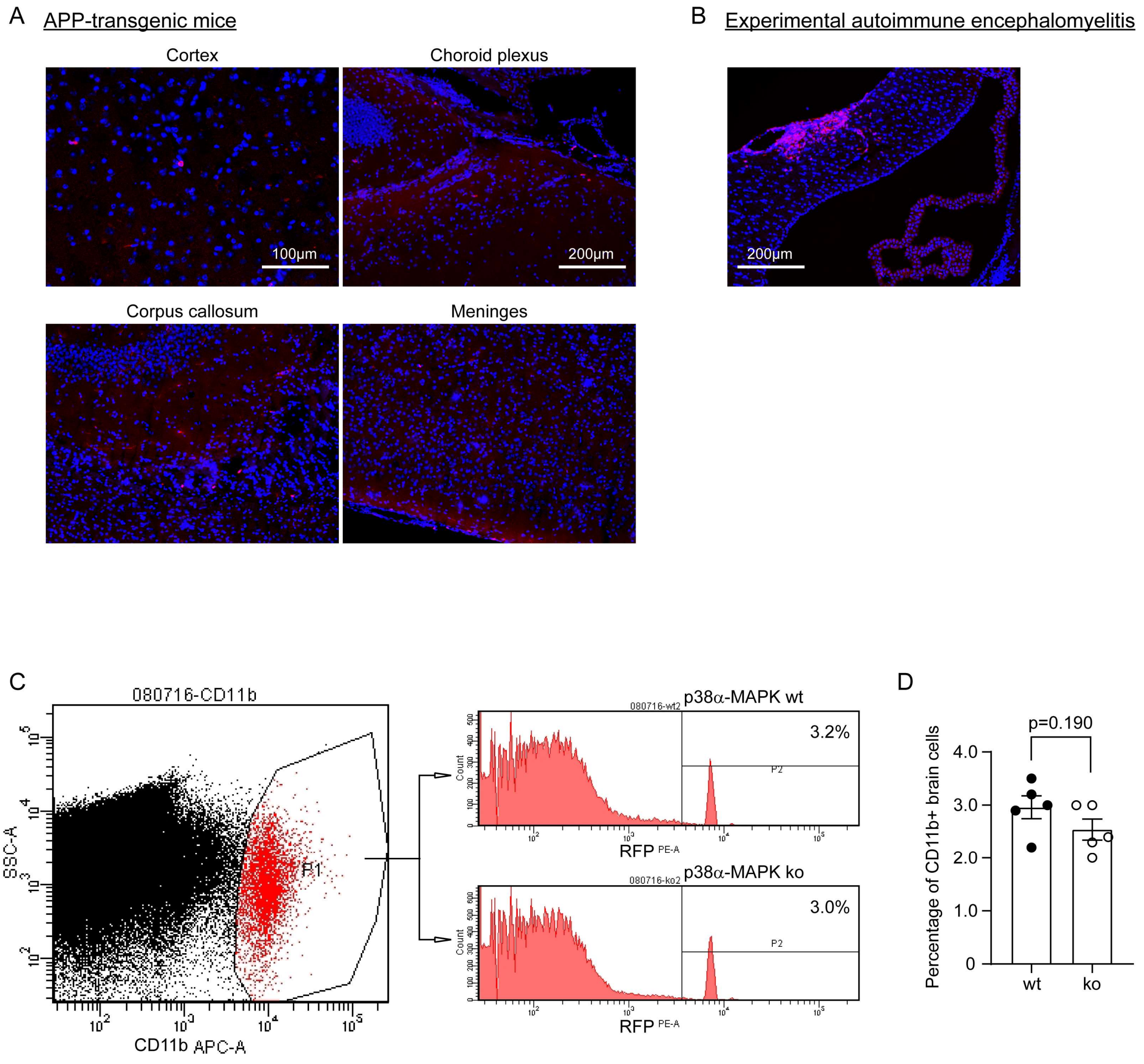


**Supplementary Fig. 1, Few peripheral myeloid cells migrate into the brain of APP-transgenic mice.** APP$^{tg}$p38$^{fl/fl}$LysM-Cre$^{+/-}$ mice were cross-bred with CCR2-RFP knock-in mice, which express RFP under the control of *ccr2* gene promoter. Paraffin-embedded brain tissues from 9-month-old APP$^{tg}$p38$^{fl/fl}$LysM-Cre$^{+/-}$CCR2$^{RFP/wt}$ mice were stained with RFP antibody (Cat.-No.: 600-401-379; Rockland Immunochemicals, Inc) and Cy3-conjugated anti-rabbit IgG. RFP-immunoreactive cells (in red) with a limited number distribute in different brain regions with a close relationship to blood vessels (A). Brain tissues from experimental autoimmune encephalomyelitis models established on CCR2$^{RFP/wt}$ mice were used as a positive control, which shows substantial RFP-positive cells clustering around blood vessels and infiltrating into the brain parenchyma (B). Single cell suspensions prepared from brains of APP$^{tg}$p38$^{fl/fl}$LysM-Cre$^{+/-}$CCR2$^{RFP/wt}$ and APP$^{tg}$p38$^{fl/fl}$LysM-Cre$^{-/-}$CCR2$^{RFP/wt}$ mice were stained with APC-conjugated CD11b antibody and analyzed with flow cytometry for RFP-expressing cells with our established protocol (Liu et al., 2012; https://doi.org/10.1016/j.neurobiolaging.2012.10.015) (C). We observed that there are 2.5% ~ 3.0% of CD11b+ brain cells in APP-transgenic mice potentially derived from peripheral myeloid cells. Deficiency of p38α-MAPK in myeloid cells does not affect the recruitment of peripheral myeloid cells into the brain, as the percentages of CCR2-RFP-positive cells among CD11b-positive brain cells were not significantly different between p38α-MAPK-deficient and wildtype APP-transgenic mice (D; t test; *n* = 5 per group).

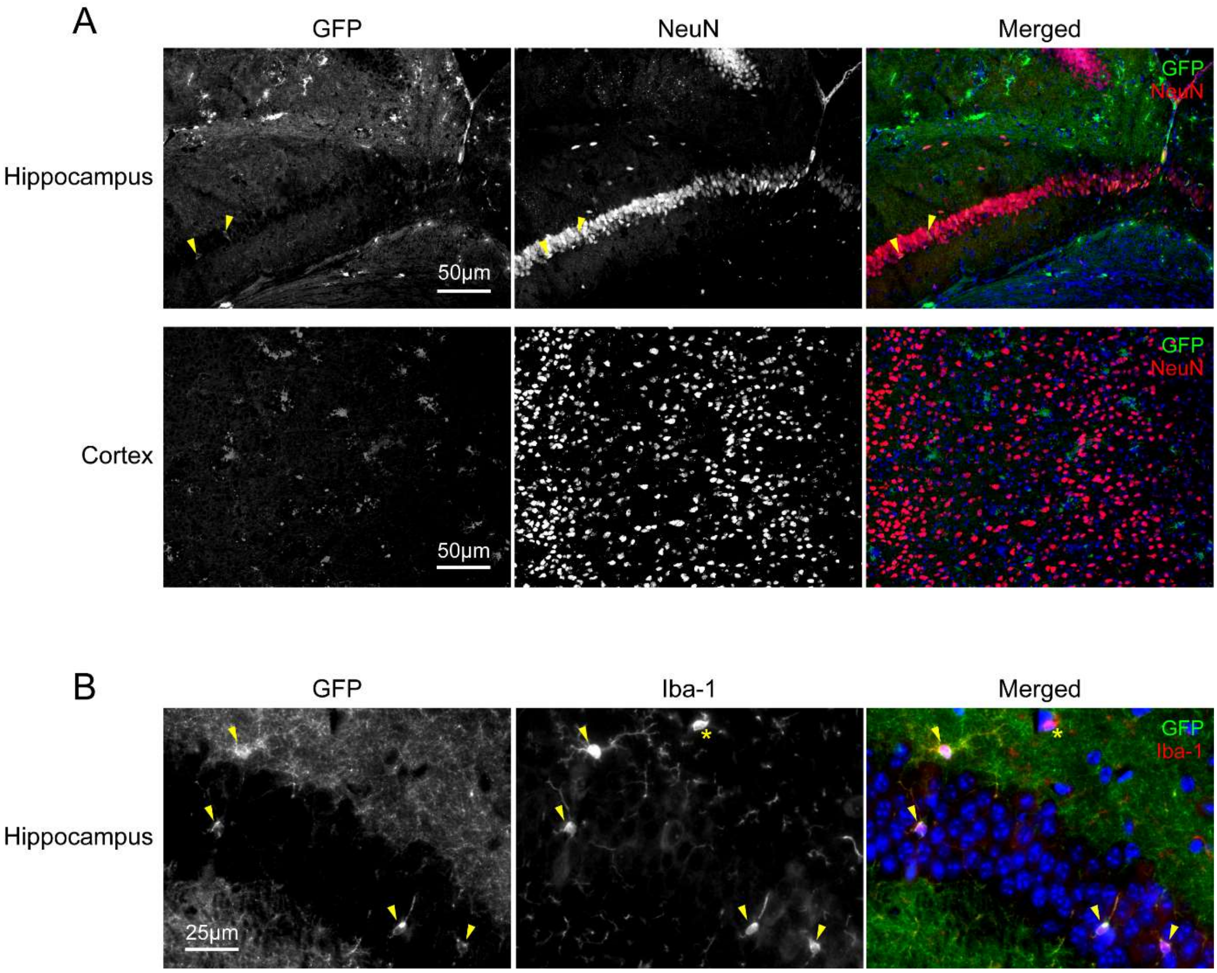


**Supplementary Fig. 2, LysM-Cre drives expression of GFP reporter rarely in neurons.** $APP^{tg}p38^{fl/fl}LysM\text{-}Cre^{+/-}$ mice were mated to $ROSA^{mT/mG}$ Cre reporter mice, which express GFP in Cre-expressing cells. Paraffin-embedded brain tissues from 6-month-old $APP^{tg}p38^{fl/wt}LysM\text{-}Cre^{+/-}ROSA^{mT/mG}$ mice were co-stained with fluorescence-conjugated antibodies against GFP (Cat.-No.: 600-401-215; Rockland Immunochemicals, Inc) and neuronal marker, NeuN (clone A60; Merck Chemicals GmbH), or microglial marker, Iba-1 (clone 20A12.1; Merck Chemicals GmbH). We observed that there are few NeuN-immunoreactive cells in the hippocampus, expressing GFP (marked with arrow heads); and there are no NeuN-positive cells in the cortex, which are positive for GFP staining (A). GFP-expressing cells in the hippocampus were clearly stained by Iba-1 antibodies (marked with arrow heads). Some Iba-1-immunoreactive cells are negative for the staining of GFP (marked with star) (B).

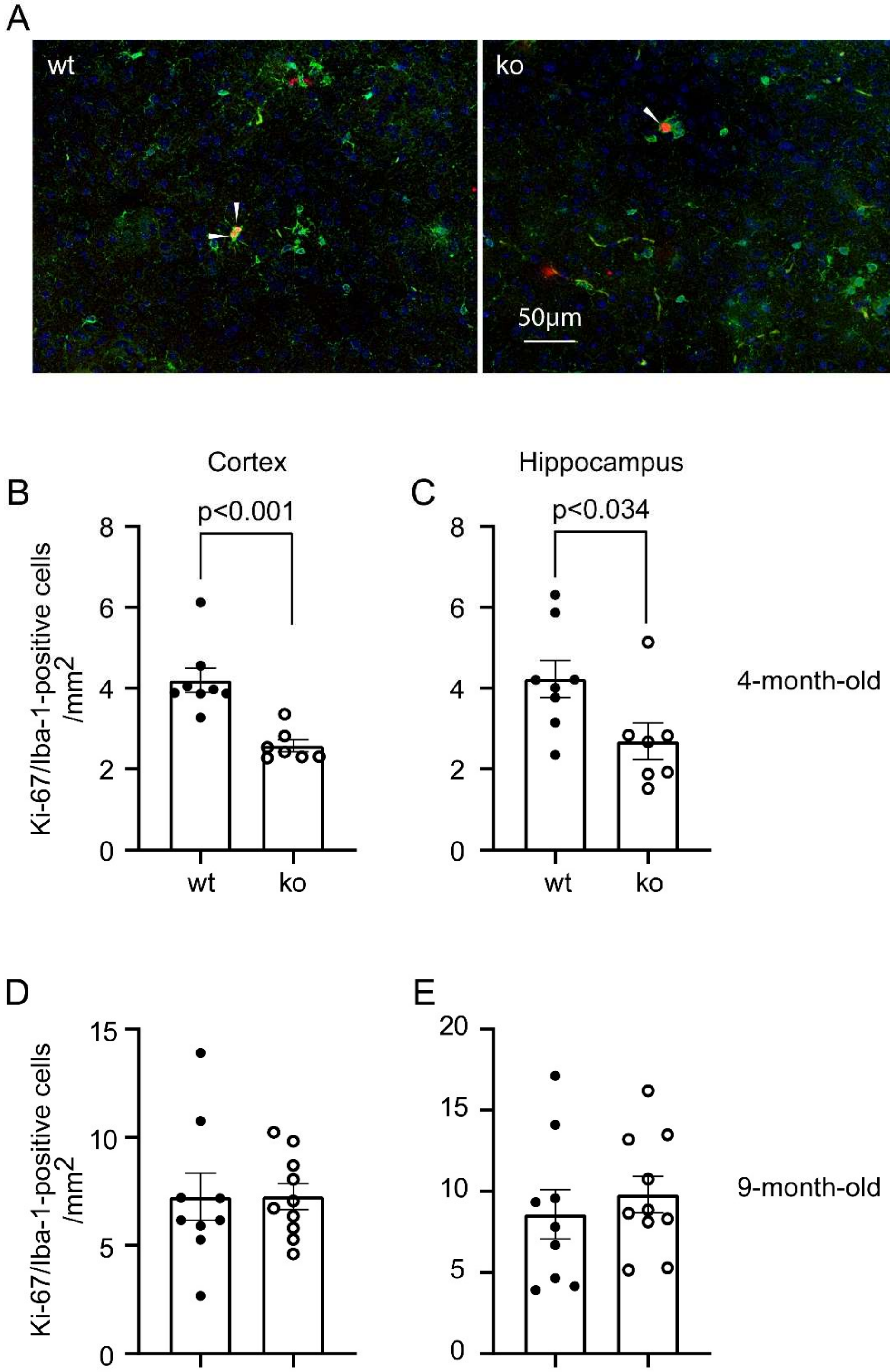


**Supplementary Fig. 3, Deletion of p38α-MAPK in myeloid cells reduces microglial proliferation in APP-transgenic mouse brain.** Four or nine-month-old APP-transgenic mice with (p38α-ko) and without (p38α-wt) deletion of p38α-MAPK in the myeloid cell lineage were analyzed for the microglial proliferation. Microglia were co-stained with fluorescence-conjugated antibodies against Iba-1 (in green; Wako Chemicals) and Ki-67 (in red; clone SP6; Abcam). Ki-67 and Iba-1-double immunoreactive cells are labeled with arrow heads (A, staining in 4-month-old APP mice). Deficiency of p38α-MAPK reduced Ki-67/Iba-1-positive cells in 4 but not in 9-month-old APP-transgenic mice (B-E; t test; $n \geq 7$ per group).

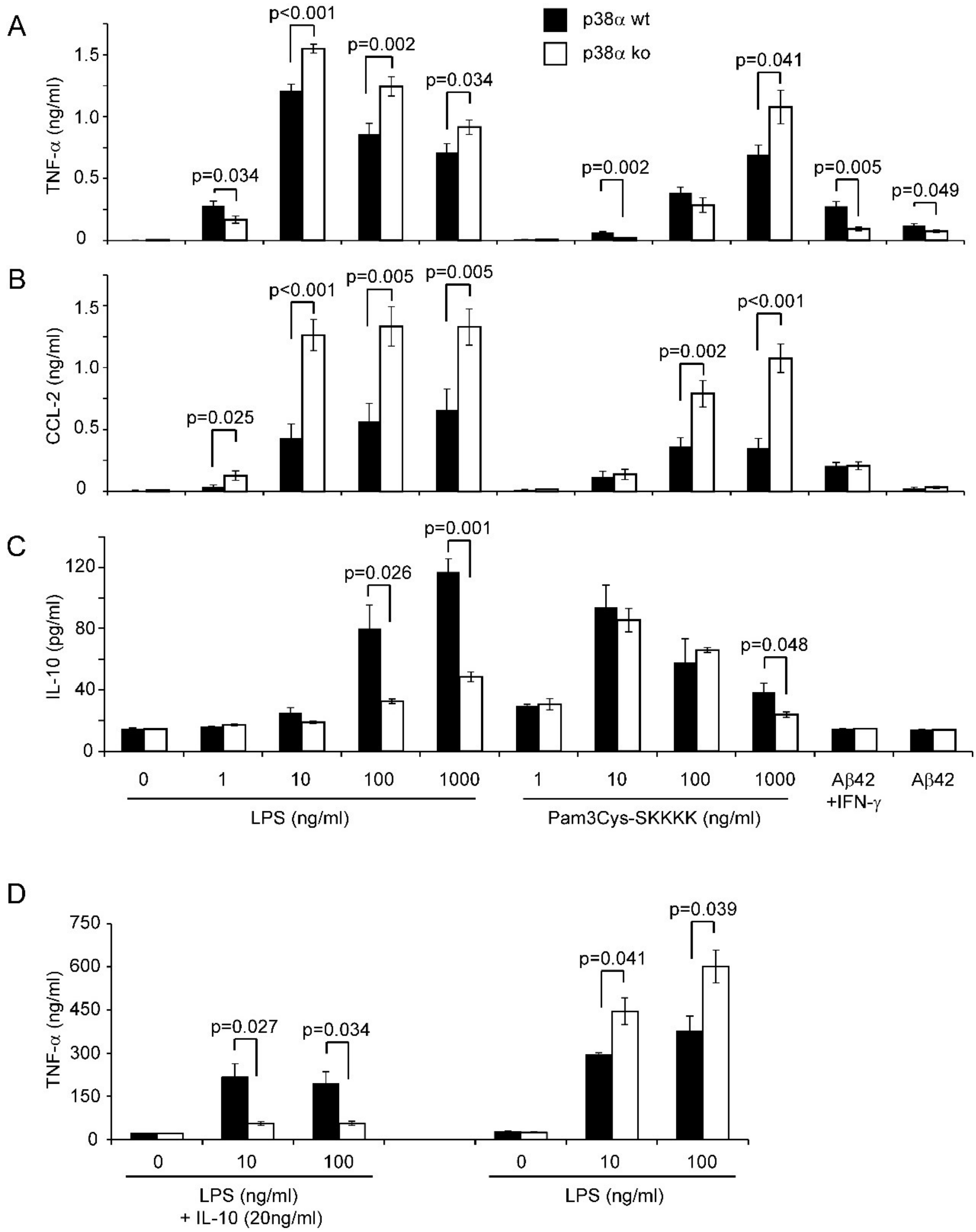


**Supplementary Fig. 4, Regulation of TNF-α-release by p38α-MAPK deficiency depends on the intensity of inflammatory stimuli.** Bone marrow-derived macrophages in 48-well plate at a density of 2 × $10^5$ cells per well were cultured from p38α-MAPK-deficient (ko) and wild-type (wt) mice with our established protocol (Hao et al., 2011; https://doi.org/10.1093/brain/awq325), and stimulated for 24 hours with LPS or Pam3Cys-SKKKK at different concentrations, or 10μM oligomeric Aβ42 aggregates in presence and absence of 100U/ml IFN-γ. Concentrations of TNF-α, CCL-2, and IL-10 in the culture medium were measured with ELISA kits (R&D Systems) (A - C; t test, $n \geq 10$ per group). p38α-MAPK wt and ko macrophages were also pre-treated with 20ng/ml mouse recombinant IL-10 (R&D Systems) for 1 hour and then stimulated with LPS in the presence of IL-10 for 24 hours (D). Co-treatment of IL-10 abolishes p38α-MAPK ko-mediated increase of TNF-α secretion (t test, $n \geq 3$ per group).

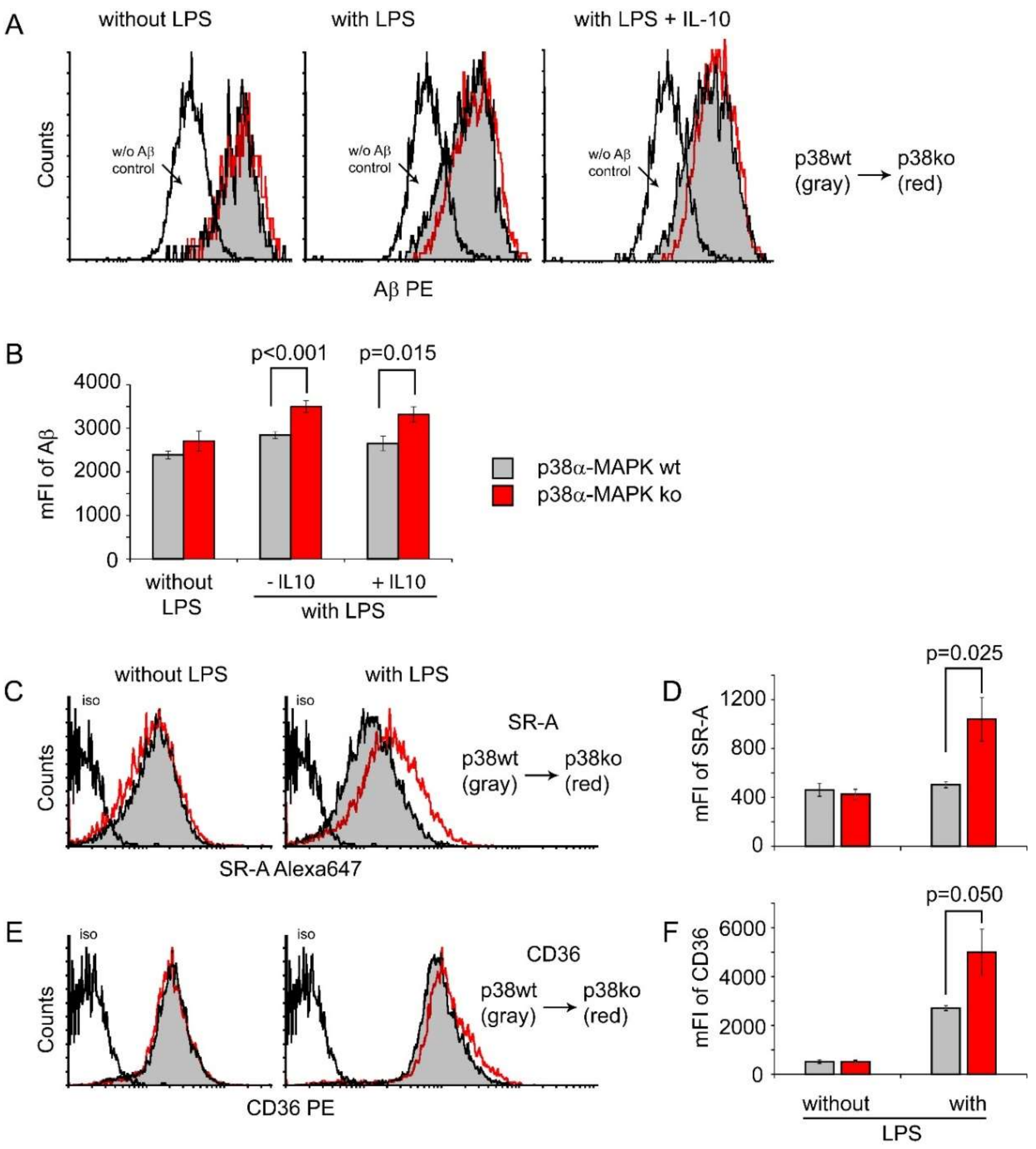


**Supplementary Fig. 5, Deletion of p38α-MAPK facilitates Aβ internalization in LPS-primed macrophages.** Bone marrow-derived macrophages in 24-well plate (BD Biosciences) at a density of $3 \times 10^5$ cells per well were cultured from $p38^{fl/fl}$LysM-$Cre^{+/-}$ (ko) and $p38^{fl/fl}$LysM-$Cre^{-/-}$ (wt) mice. They were primed with and without 100ng/ml LPS for 48 hours. Some cells were co-treated with recombinant mouse IL-10 at 20ng/ml. Fluorescent Aβ aggregates were prepared by mixing TAMRA-labeled human Aβ42 (Anaspec Inc.) and unlabeled Aβ42 (kindly provided by L. Fülöp, Albert Szent Gyorgyi Medical University, Szeged, Hungary) at a ratio of 1:10 and incubating 100µM mixed peptides in phenol red-free Ham's F-12 for 24 hours, as we did in a previous study (Liu et al., 2014; https://doi.org/10.1523/JNEUROSCI.1348-14.2014). Thereafter, macrophages were fed with 10µM TAMRA-conjugated oligomeric Aβ42 in the presence of LPS with and without IL-10 for 18 hours. The internalization of Aβ was monitored by measuring mean fluorescence intensity (mFI) with flow cytometry (A and B; *t* test, $n \geq 6$ per group). Moreover, protein levels of SR-A and CD36 were detected with flow cytometry after immunofluorescent staining of macrophages with Alexa647-labelled rat anti-mouse SR-A (clone 2F8; Bio-Red), and PE-labeled rat anti-mouse CD36 (clone HM36; Thermo Fisher Scientific), respectively (C - F; *t* test, $n \geq 4$ per group).

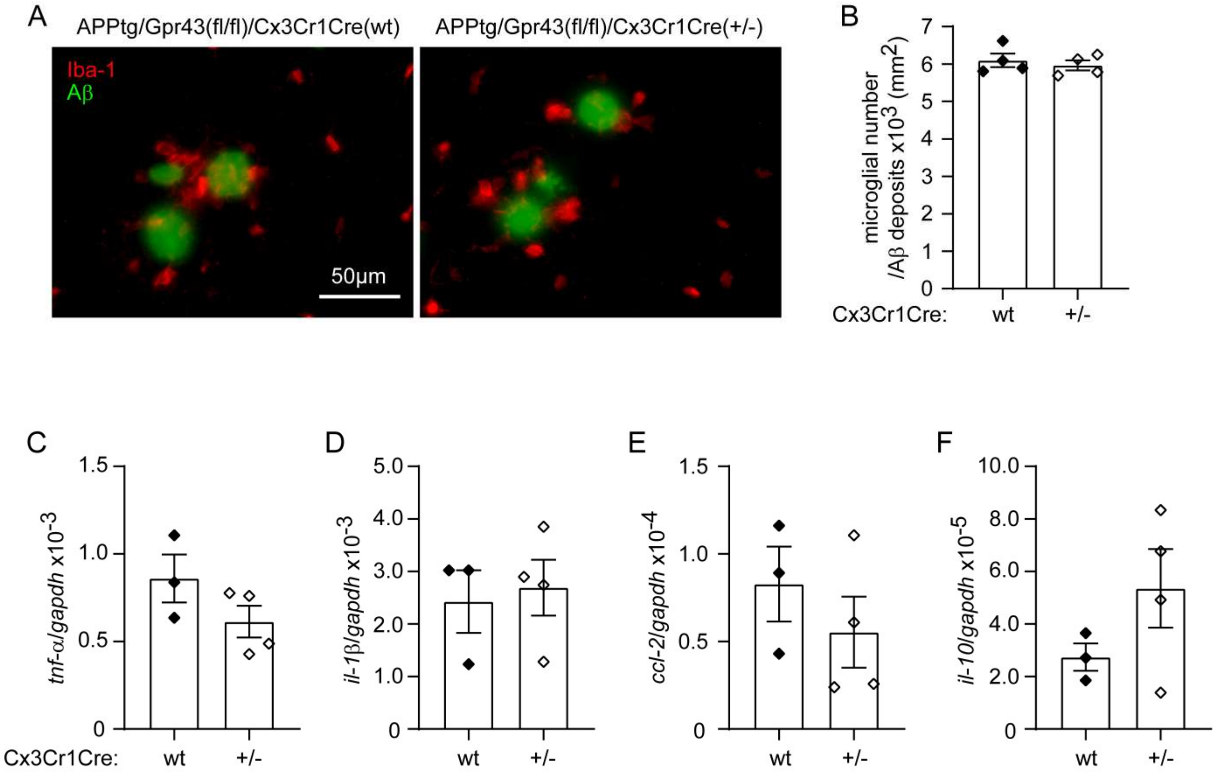


**Supplementary Fig. 6, Haploinsufficient Cx3Cr1 affects microglia for neither migration to Aβ deposits nor inflammatory gene transcription in APP-transgenic mice.** APP/PS1-transgenic mice were cross-bred with Cx3Cr1-CreERT2 mice and *gpr43*-floxed mice (Tang et al., 2015; https://doi.org/10.1038/nm.3779) to obtain mice with APP$^{tg}$Gpr43$^{fl/fl}$Cx3Cr1-Cre$^{+/-}$ and APP$^{tg}$Gpr43$^{fl/fl}$Cx3Cr1-Cre$^{-/-}$ of genotypes. These two groups of littermate mice were injected (*i.p.*) with tamoxifen and analyzed for AD-associated pathology by 9 months. As GPR43 is not expressed in microglia (Quan et al., 2021; https://doi.org/10.1002/glia.24007), the pathological difference between APP$^{tg}$Gpr43$^{fl/fl}$Cx3Cr1-Cre$^{+/-}$ and APP$^{tg}$Gpr43$^{fl/fl}$Cx3Cr1-Cre$^{-/-}$ mice should come from the haploinsufficiency of Cx3Cr1. In recent studies, we observed that haploinsufficiency of Cx3Cr1 changes neither Aβ load nor neuroinflammation (Quan et al., 2021). In this study, we further co-stained brain tissues with Cy3-conjugated antibodies against Iba-1 (clone E4O4W; Cell Signaling Technology) and methoxy-XO4 (Bio-Techne GmbH), and demonstrated that haploinsufficiency of Cx3Cr1 does not change microglial recruitment to Aβ deposits (A and B; *t* test, $p > 0.05$, $n = 4$ per group). Moreover, we selected CD11b-positive cells from 9-month-old APP$^{tg}$p38$^{fl/fl}$Cx3Cr1-Cre$^{+/-}$ and APP$^{tg}$p38$^{fl/fl}$Cx3Cr1-Cre$^{-/-}$ littermate mice, which had not been treated with tamoxifen for the induction of gene recombination. As detected with real-time PCR, we observed that haploinsufficiency of Cx3Cr1 does not alter the transcription of inflammatory genes, *tnf-α*, *il-1β*, *ccl-2* and *il-10* (C - F; *t* test, $p > 0.05$, $n \geq 4$ per group).

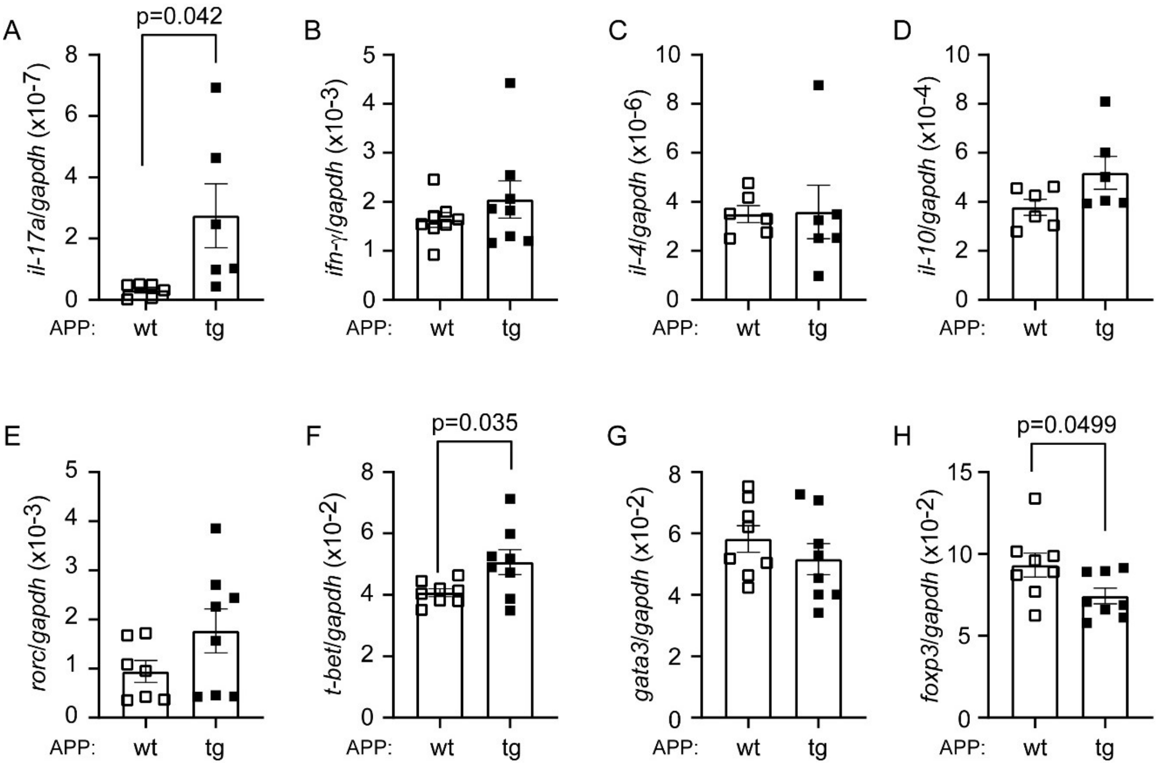


**Supplementary Fig. 7, Transcription of *il-17a* is up-regulated in CD4+ spleen cells in APP-transgenic mice.** CD4-positive spleen cells were selected with magnetic beads-conjugated antibodies from 6-month-old APP-transgenic and wildtype littermate mice and detected with real-time PCR for transcription of T lymphocyte marker genes. The transcription of *il-17a*, but not *ifn-γ*, *il-4* and *il-10* genes is significantly up-regulated in CD4+ spleen cells of APP-transgenic mice, compared with APP-wildtype littermates (A - D). *t* test; $n \geq 6$ per group).
